# Live-cell imaging and ultrastructure expansion microscopy reveal the dynamic intracellular life cycle of the microsporidian parasite *Encephalitozoon intestinalis*

**DOI:** 10.64898/2026.09.25.754474

**Authors:** Kacie McCarty, Sabrina Absalon, Gira Bhabha, Damian Ekiert

## Abstract

Microsporidia are divergent fungal pathogens that infect a wide range of hosts. Microsporidia have evolved highly reduced genomes, resulting in the loss of many metabolic pathways. *Encephalitozoon intestinalis* is a human-infecting microsporidian species with the smallest known eukaryotic genome (2.3 Mbp). As an obligate intracellular parasite, *E. intestinalis* is dependent on its host for replication. Our current understanding of *E. intestinalis* cell biology remains limited. This is largely due to genetic intractability, which has made live-cell imaging challenging, and its small size (∼2 µm), which is within the diffraction limit of conventional microscopy. Here we develop a live-cell imaging technique, “silhouette microscopy”, that does not rely on genetic manipulation of microsporidia, and enables us to study the live cell dynamics of *E. intestinalis* replication in host cells, from entry to exit. We report the timing of cell division, kinetics of the parasite life cycle, and dynamics of the parasitophorous vacuole within which *E. intestinalis* resides. The parasitophorous vacuole undergoes frequent fission and fusion, and ultimately ruptures to facilitate parasite release from the host cell. Silhouette microscopy provides an excellent platform to study the life cycle dynamics of other microsporidian species as well as other genetically intractable intracellular pathogens. To gain a higher resolution understanding into the cell biology of *E. intestinalis* development, we performed ultrastructure expansion microscopy (U-ExM) on infected cells. U-ExM reveals the organization of the *E. intestinalis* cytoskeleton and development of the specialized invasion organelle, the polar tube. The combination of live-cell imaging and U-ExM provide a platform to study the *E. intestinalis* life cycle, and sheds light on the dynamics and cell biology of *E. intestinalis* in host cells.

## Introduction

Microsporidia are an early-diverging group of fungal pathogens that infect a wide range of invertebrate and vertebrate hosts (1–4). In humans, microsporidia infection is frequently caused by the species *Encephalitozoon intestinalis*, which primarily infects the small intestine (5, 6). *E. intestinalis* infection typically results in self-limiting diarrheal disease. However, in immunocompromised individuals, disseminated disease has been observed and can be fatal (7, 8). *E. intestinalis* is transmitted between hosts in the form of environmentally resistant, dormant spores, the only stage of the life cycle that can survive outside a host. Entry into a host cell is mediated by a unique harpoon-like organelle, called the polar tube (PT) (9, 10). The PT, which is several times longer than the spore itself, is initially coiled within the dormant spore (11). Once triggered, the PT rapidly extends from the spore body, transforming from a coil to a linear tube. This hollow tube serves as a conduit for the transport of infectious cargo (also called sporoplasm) from the spore body into the host cell (9). Upon entry into the host cell, *E. intestinalis* replicates within a parasitophorous vacuole (PV) and remains within the PV throughout the life cycle. As an obligate intracellular parasite, *E. intestinalis* has co-evolved with, and is dependent upon, the host for replication and the completion of its life cycle (12–14).

Following entry of the sporoplasm, the *E. intestinalis* life cycle is characterized by a series of four morphologically distinct stages (**Fig. 1A**). The life cycle begins at the proliferative stage, in which *E. intestinalis* cells replicate via binary fission (4). After several rounds of replication, the proliferative cells transition to the sporont stage where abscission of septated parasites is completed and the spore coat starts to form (4, 15–17). Shortly thereafter, *E. intestinalis* enters the sporoblast stage in which specialized organelles, such as the PT, begin to develop (4, 15–17). The final stage of development is the spore stage. Mature spores are surrounded by a thick spore wall, composed of a proteinaceous exospore layer and a polysaccharide-based endospore layer, which includes chitin and β-glucans (4, 15–17).

**Figure 1.**
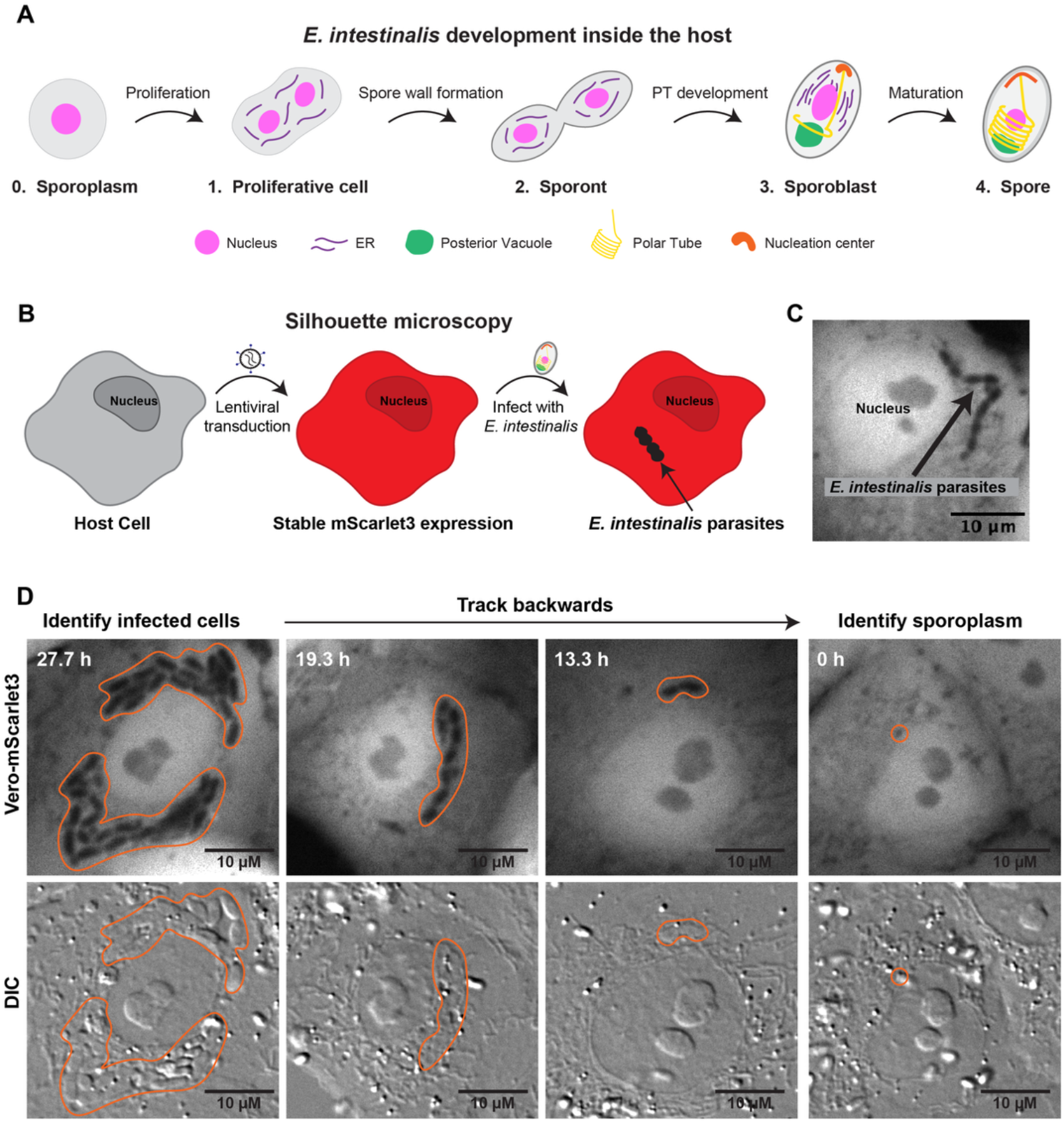
Silhouette microscopy as a tool to study live-cell dynamics of the *E. intestinalis* life cycle. A. Schematic representing our current understanding of the *E. intestinalis* life cycle, which occurs inside host cells. The sporoplasm (0) enters the host cell, after which *E. intestinalis* goes through four morphologically distinct life stages: replication occurs at the proliferative cell stage (1), followed by two sequential developmental stages: sporont (2) and sporoblast (3). Finally, mature spores (4) are formed and released from the host cell. **B**. Schematic overview of the silhouette microscopy experimental workflow. Host cells are transduced with a lentivirus to stably express mScarlet3. Cells expressing mScarlet3 are then infected with *E. intestinalis* and immediately imaged. When infected, the parasites remain dark against the bright fluorescent background of the host cell creating a “silhouette”. **C**. Representative image of a Vero-mScarlet3 cell that is infected with *E. intestinalis*, which appear as dark silhouettes against the bright host cell cytoplasm. **D**. Frames of a movie capturing an *E. intestinalis* infection event in fluorescence and DIC channels. Infected cells are easily identified in later frames of each movie in both channels, after replication. The movie is tracked backwards to capture the entry of a single silhouette in an earlier frame, which represents the sporoplasm in the host cell. *E. intestinalis* parasites are outlined in orange in each frame, which are visible in the fluorescence channel, but not in DIC.

Much of our knowledge regarding the microsporidian life cycle comes from fixed imaging techniques such as electron microscopy (18) and immunofluorescence (19). These imaging techniques have been instrumental in laying the groundwork for our understanding of *E. intestinalis* development, but are limited in throughput and resolution, respectively. In addition, neither electron microscopy nor immunofluorescence capture the live-cell dynamics of parasite replication. As such, we are missing key information into the life cycle such as the timing of cell division, kinetics of the parasite life cycle, and dynamics of the PV. This is largely due to the genetic intractability of microsporidia, which has rendered fluorescent tags for live-cell imaging unfeasible. The small size of *E. intestinalis* also limits our ability to study parasite development and subcellular organization, and requires high resolution microscopy techniques to visualize cellular features of individual parasite cells.

To facilitate live-cell imaging in microsporidia, we developed a technique that does not rely on genetic manipulation of microsporidia, and provides single parasite-level resolution, which we term “silhouette microscopy”. Using silhouette microscopy, we report on the entire *E. intestinalis* life cycle in live host cells, from entry to exit, including parasite replication and PV dynamics. In addition, to obtain higher resolution information on the cell biology of *E. intestinalis* through its developmental stages, we turned to ultrastructure expansion microscopy (U-ExM). We adapted a U-ExM protocol (20, 21) to establish expansion microscopy as a robust technique for studying microsporidia. This provides the throughput and labeling-specificity of immunofluorescence, at resolutions approaching those achieved by volume electron microscopy imaging of *E. intestinalis-*infected host cells (17). Using U-ExM, we gained an understanding into the organization of the *E. intestinalis* cytoskeleton and the development of the polar tube through the life cycle. Combining silhouette microscopy and U-ExM, we present a temporal framework for *E. intestinalis* replication from entry to exit, and snapshots of *E. intestinalis* development at ∼40-50 nanometer resolution.

## Results

### Developing silhouette microscopy to study microsporidia

To characterize the dynamics of *E. intestinalis* infection, we set out to perform live-cell imaging of infected cells. To overcome the lack of tools for live-cell imaging in microsporidia, we generated mammalian cell lines that overexpress cytoplasmic mScarlet3. When infected, the mScarlet3 creates a fluorescent background in the host cell while the parasites remain dark, creating a “silhouette” (**Fig. 1B and C**). We have termed this imaging technique “silhouette microscopy”, and it enables us to visualize the *E. intestinalis* life cycle in living cells. We performed silhouette microscopy in Vero-mScarlet3 cells, a monkey kidney epithelial cell line often used to propagate *E. intestinalis* spores, and Caco-2-mScarlet3 cells, a colonic cell line which provides a more physiologic environment to study parasite dynamics, and has previously been used as a model cell line (22). Based on fixed imaging analysis, a similar timeline for the *E. intestinalis* life cycle in Vero and Caco-2 cells was observed (23). Using these infection timelines as a guide, we chose a time window for live-cell silhouette imaging. To capture the entire *E. intestinalis* life cycle, we began imaging immediately after infection and continued to image for ∼65 h. During the early stages of parasite replication, the parasites cannot be seen via DIC, due to their flattened appearance, but can be observed by silhouette microscopy (**Fig. 1D**). At the start of imaging, fields of view were selected at random, as host cells are not yet infected. To analyze the parasite life cycle, later frames of each movie, in which infected cells are clearly visible, were identified first. Using these frames as a starting point, the movie was tracked backwards to capture the entry of a single sporoplasm, which appears as a single silhouette (**Fig. 1D, Movie S1**). Using silhouette microscopy we can track the entire parasitic life cycle in live cells from entry to exit.

### Replication kinetics of *E. intestinalis* using silhouette microscopy

To gain a better understanding of parasite replication kinetics, we monitored *E. intestinalis* division beginning immediately after entry via silhouette microscopy and corresponding DIC imaging in both Vero-mScarlet3 and Caco-2-mScarlet3 cells. Vero cells grow in flat monolayers and are generally featureless except for the silhouettes, making live-cell imaging robust, and the data of high quality. Caco-2 cells, on the other hand, tend to grow in clusters rather than an even monolayer, and often exhibit numerous dark vesicle-like structures in the cytoplasm (**Fig. S1**). This makes live-cell silhouette imaging of *E. intestinalis* in Caco-2 cells more challenging. For these reasons, we collected and quantified more infection events in Vero cells than in Caco-2 cells. Overall, our findings are similar in both host cell lines. Upon entry of the sporoplasm, the parasites undergo a lag time of ∼15.5 +/-2.7 h in Vero cells and ∼13.9 +/-1.8 h in Caco-2 cells before completion of the first cell division (**Fig. 2A, C, F, and H, Movies S2 and S3, and Datasets S1 and S2**). This observation is consistent with the lag time observed for the microsporidian species, *Nematocida parisii*, which was also reported to undergo an ∼18 h lag phase when infecting *Caenorhabditis elegans* (24), based on analysis of fixed cells. The lag phase suggests that *E. intestinalis, N. parisii* and possibly other microsporidia may require significant time post-invasion to initiate gene expression, transition from a dormant to actively replicating state, and perhaps repair genomic damage following the extreme nuclear deformation that occurs during translocation through the PT (11). Single-cell RNA sequencing data of *E. intestinalis* replicating within macrophages is also consistent with a lag time post invasion at the transcriptional level, suggesting that the parasites may indeed require time to initiate gene expression over several hours following host entry (25).

**Figure 2.**
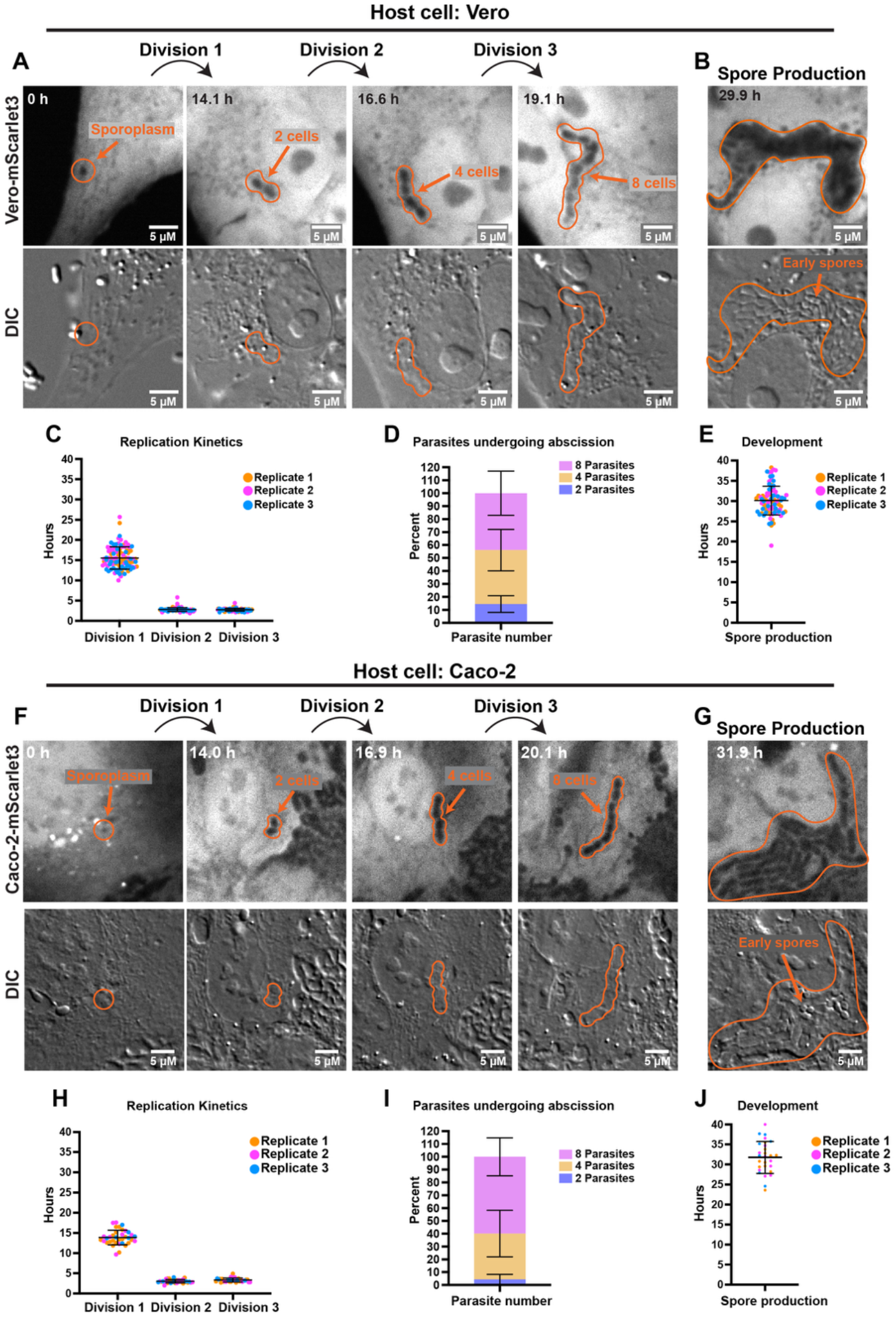
Replication kinetics of *E. intestinalis* in Vero and Caco-2 host cells. A, B. Vero-mScarlet3 cells infected with *E. intestinalis* and live-cell imaging was performed from the time of infection. Frames of a movie are shown, using silhouette microscopy (top panel) and DIC (bottom panel) to track parasite division (**A**) and spore production (**B**). Parasite boundaries are marked with solid orange outlines. **C**. Quantification of the time for the first three parasite divisions after entry of the sporoplasm into Vero-mScarlet3 cells. Quantification is from three replicates, and error bars the SD. Replicate 1, *n* = 29 parasites; Replicate 2, *n* = 33 parasites; Replicate 3, *n* = 34 parasites. **D**. Quantification of the number of parasites undergoing the first abscission at the 2, 4, and 8 parasite cell stages in Vero-mScarlet3 cells. Quantification is from three replicates, and error bars represent the SD. *n* = 83 parasites were analyzed. **E**. Quantification of the timing of spore production in Vero-mScarlet3 cells, based on the transition in contrast of DIC imaging. Quantification is from three replicates, and error bars represent the SD across the three replicates. *n* = 86 parasites were analyzed. **F, G**. Caco-2-mScarlet3 cells infected with *E. intestinalis* and live-cell imaging was performed from the time of infection. Frames of a movie are shown, using silhouette microscopy (top panel) and DIC (bottom panel) to track parasite division (**F**) and spore production (**G**). Parasite boundaries are marked with solid orange outlines. **H**. Quantification of the time for the first three parasite divisions after entry of the sporoplasm into Caco-2-mScarlet3 cells. Quantification is from three replicates, and error bars represent the SD. Replicate 1, *n* = 18 parasites; Replicate 2, *n* = 11 parasites; Replicate 3, *n* = 7 parasites. **I**. Quantification of the number of parasites undergoing the first abscission at the 2, 4, and 8 parasite cell stages in Caco-2-mScarlet3 cells. Quantification is from three replicates, and error bars represent the SD. *n* = 35 parasites were analyzed. **J**. Quantification of the timing of spore production in Caco-2-mScarlet3 cells, based on the transition in contrast of DIC imaging. Quantification is from three replicates, and error bars represent the SD across the three replicates. *n* = 28 parasites were analyzed.

Following the lag time, *E. intestinalis* replicates via binary fission, and each division takes ∼2.7 +/-0.4 h in Vero cells and ∼3.2 +/-0.5 h in Caco-2 cells (**Fig. 2A, C, F, and H, Movies S2 and S3, and Datasets S1 and S2**). After each round of nuclear replication, 2 daughter cells are formed and undergo furrow ingression. However, abscission is not completed, resulting in chains of connected cells, resembling sausage links. The first division yields two connected cells, followed by four and then eight (**Fig. 2A and F**). The chains grow to a maximum of eight cells, which ultimately undergo asynchronous abscission. The chains of cells most often undergo abscission at the four and eight cell stages, but can also undergo abscission at the two cell stage (**Fig. 2D and I, Fig. S2A and B**). After abscission of the initial chains, the resulting parasites continue to replicate and divide, before beginning to differentiate into spores. While replicative and early developing parasites are not visible using DIC alone, later in infection, parasites become clearly visible by DIC, perhaps due to shrinkage and a change in contrast during spore development. To assess if parasites visible by DIC indeed represent maturing spores, we fixed and stained infected cultures with an RNA fluorescence in situ hybridization (FISH) probe against *E. intestinalis* 16S rRNA, and calcofluor white, which labels chitin. Under our conditions, the RNA FISH probe stains immature parasites, but is excluded from mature parasites, as the spore wall blocks entry of the FISH probe. Calcofluor white exclusively stains mature parasites that have begun to develop their cell wall. Analysis of fluorescence and corresponding DIC images shows that 100% of parasites that are clearly visible by DIC are also chitin positive, and are RNA FISH negative, suggesting that these are maturing parasites (**Fig. S3**). We can therefore use the transition we observe in DIC imaging to assess parasite maturity in live cells, and we refer to these clearly visible parasites as “maturing parasites”. Maturing parasites begin to appear at approximately ∼30.1 +/-3.5 hpi in Vero cells and ∼31.8 +/-4.6 hpi in Caco-2 cells (**Fig. 2B, E, G, and J and Movie S2 and S3**). While some parasites begin to mature at these timepoints, many other parasites within the same host cell continue to replicate, creating an asynchronous infection within a single PV, consistent with previous observations (17). Overall our data allow us to characterize the timing and dynamics of *E. intestinalis* replication in real time.

### Ultrastructure expansion microscopy captures subcellular features of developing *E. intestinalis* parasites

Silhouette microscopy allows us to capture the dynamics of *E. intestinalis* at the level of single cells. However, as each parasite is a dark silhouette, we are unable to learn about the subcellular organization of individual parasites using this technique. Due to the small size of *E. intestinalis* (∼2 µm), standard immunofluorescence techniques are not sufficient to capture the organization of organelles and cytoskeleton of an individual parasite. To gain higher resolution insights into developing parasites with reasonable throughput, we turned to U-ExM, which enables us to achieve higher resolution information with the specificity and throughput of immunofluorescence. U-ExM results in a ∼4 fold isotropic expansion of the sample (20) and has been successfully used to study several microbial organisms such as *Plasmodium falciparum* (26–28). The physical increase in sample size, coupled with antibody and dye staining, allows for the visualization of subcellular features in small cells, making it an ideal approach for the study of microsporidia. Starting from a U-ExM protocol previously used to study *P. falciparum* (21), we adapted it to study *E. intestinalis*-infected cells. This protocol allows us to resolve subcellular features within single *E. intestinalis* parasites. For example, microtubules in mitotic spindles are clearly visible in expanded samples (**Fig. 3A**). To capture several stages of parasite development, we fixed infected Vero cells at 20, 26, and 30 hpi. We stained our samples for DNA (DRAQ5), *E. intestinalis* β-tubulin (polyclonal antibody), and general cellular features using a non-specific protein-labeling NHS-ester dye. For each infection timepoint, we acquired 3 data sets from which we captured 111 infected cells containing 154 PVs: 63 PVs from 20 hpi, 45 PVs from 26 hpi, and 46 PVs from 30 hpi.

**Figure 3.**
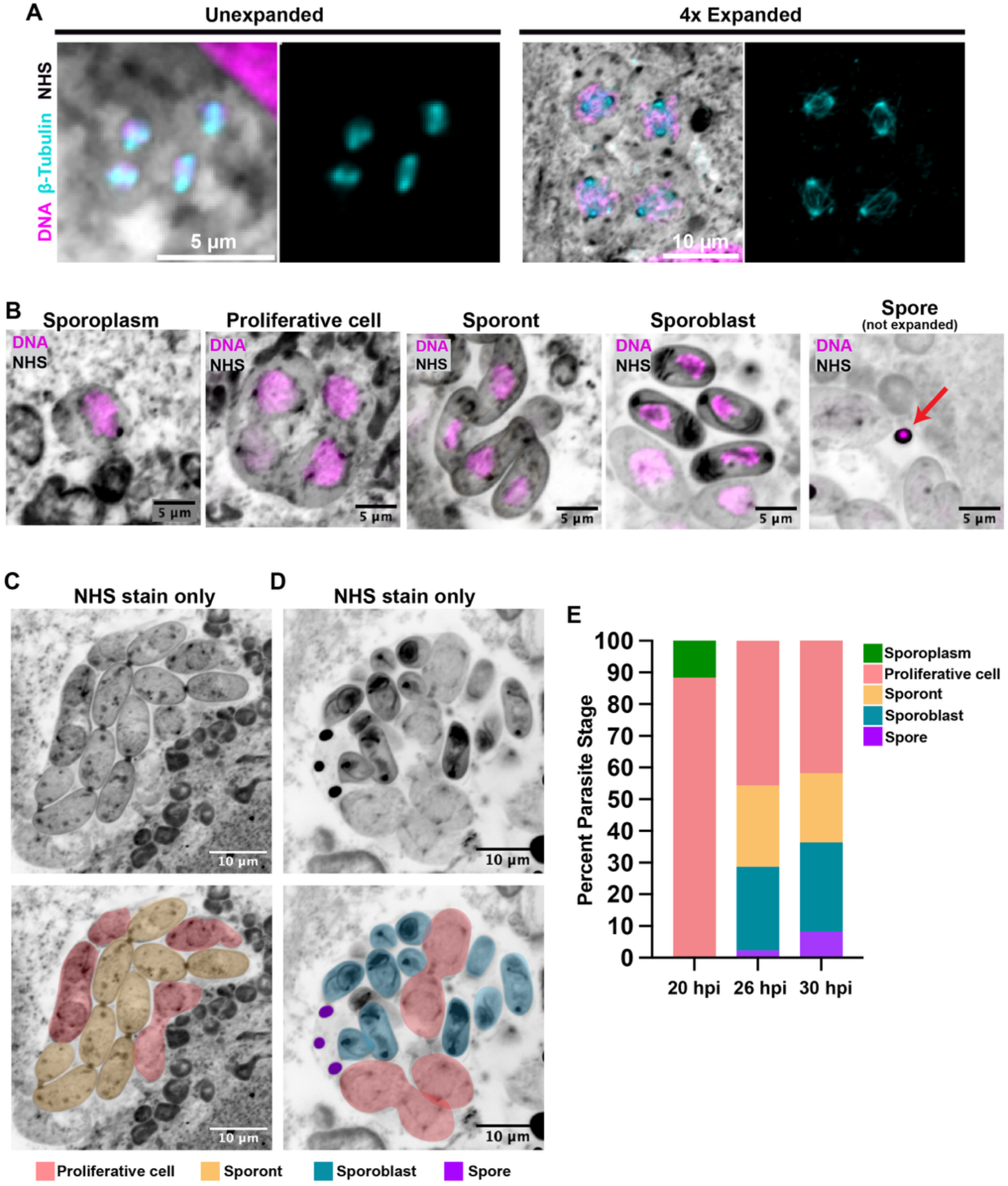
Ultrastructure Expansion Microscopy (U-ExM) of intracellular *E. intestinalis* parasites in Vero host cells. A. Comparison of unexpanded and 4x expanded Vero cells infected with *E. intestinalis*. Vero cells were infected with *E. intestinalis* for 26 h and then stained for DNA (DRAQ5, magenta), protein (NHS-ester, grayscale), and *E. intestinalis* β-tubulin (custom antibody, cyan). Individual microtubules are resolved in the 4x expanded samples prepared for U-ExM. **B**. Vero cells infected with *E. intestinalis* were prepared for U-ExM and stained for DNA (DRAQ5, magenta) and protein (NHS-ester, grayscale). Representative maximum intensity projections of each parasite developmental stage. Fully mature spores (red arrow) do not expand under our experimental conditions. **C, D**. Representative maximum intensity projections of single PVs containing parasites at different parasite developmental stages. All images were deconvolved in the Nikon NIS Elements AR software using the Blind deconvolution method (3 iterations). **E**. Quantification of the percentage of each developmental stage at each time point. Across three independent experiments *n* = 198 parasites at 20 hpi, *n* = 662 parasites at 26 hpi, and *n* = 728 parasites at 30 hpi.

Across PVs, we identified all stages of the parasite life cycle: single cell sporoplasms, chains of proliferative cells and sporonts, and individual sporoblasts and spores (**Fig. 3B**). Due to the thick chitin-containing spore wall, mature spores do not expand under our conditions, and likely will require adapted protocols to disrupt the spore wall (29). Therefore, while we can identify spores in our data set as smaller, protein dense bodies containing DNA, we are missing subcellular information of the fully mature spores (**Fig. 3B**). Consistent with our silhouette microscopy data and previous electron microscopy data, parasite development is asynchronous. A single PV contains parasites at different developmental stages (**Fig. 3C and D**), and the proportions of the parasite states changes over the course of infection (**Fig. 3E**). At 20 hpi, PVs consist of only sporoplasms or proliferative cells. At 26 hpi, sporoplasms are not observed, the percentage of proliferative cells decreases, and later developmental stages appear. At 30 hpi, a shift towards later developmental stages is observed. These observations align with our silhouette microscopy data, in which we typically observe ∼4-8 proliferative parasites per infected cell at 20 hpi. By 30 hpi, maturing parasites are apparent by DIC in live-cell imaging, in line with the shift towards sporoblasts and spores observed in our U-ExM data.

### Dynamics of the *E. intestinalis* cytoskeleton

Microtubules are an essential component of the eukaryotic cytoskeleton, playing critical roles in cell division, maintenance of cellular architecture, and cellular motility (30). Although cytoskeletal components are conserved across eukaryotes, cytoskeletal organization across organisms varies widely (26). How the cytoskeleton is organized and remodeled throughout the *E. intestinalis* life cycle remains poorly understood. Previous transmission electron microscopy data identified mitotic spindles in *E. hellem*, a related human-infecting species of microsporidia. These transmission electron microscopy data also suggest that *E. hellem* proliferates through multiple rounds of nuclear division without disassembly of the nuclear envelope, indicative of semi-open or closed mitosis (31). Tracking the entire cell cycle has been limited due to the low throughput of electron microscopy and lack of antibodies available targeting the cytoskeleton. Using U-ExM we are able to clearly visualize tubulin spindles in the parasites which were previously not resolvable by immunofluorescence (**Fig. 3A**). This much higher resolution imaging of tubulin than previously achievable allows us to track each stage of mitosis, the architecture of the microtubule cytoskeleton, and the spindle pole bodies (31, 32). During interphase, each parasite contains 1 spindle pole body associated with the nuclear envelope with cytoplasmic microtubules radiating from the spindle pole body (**Fig. 4A**). At the start of mitosis, the spindle pole bodies have already been duplicated and begin migrating to opposite poles, suggestive of prophase. During metaphase, intranuclear mitotic spindles form while the nuclear envelope appears to remain intact. Unlike open mitosis, the chromosomes do not line up at the metaphase plate, but instead remain as DNA clusters. During anaphase, the spindles elongate to form linear bundles and begin to separate the replicated DNA clusters. A hallmark of semi-open and closed mitosis is that the nuclear envelope takes on a dumbbell shape, which we observe by NHS staining (**Fig. 4A**). In telophase, the nuclear envelope is divided in two and the nuclei are separated at either pole, ready to initiate another round of replication. Throughout the cell cycle, parasites in the same chain remain synchronized at the same cell cycle stage (**Fig. S4**), suggesting that the asynchronicity of infection occurs post-abscission.

Interestingly, the frequency with which we observe proliferative cells and sporonts at each cell cycle stage varies. We predominantly capture proliferative cells in mitosis (∼92%), especially prophase (76%), whereas we capture most sporonts in interphase (65%) and a smaller fraction of sporonts in mitosis (35%) (**Fig. 4B**). This suggests that sporonts are capable of dividing; however, the proliferative stage is likely responsible for most parasite replication, and, upon entering the sporont stage, we observe a shift away from replication and toward differentiation into spores (**Fig. 3E and Fig. 4B**). Consistent with this observation, the spindle pole bodies are no longer duplicated and microtubules remain cytoplasmic. Towards the end of the sporont stage, the chains of sporonts complete a final round of abscission to form individual sporoblasts (**Fig. 4C**). Sporoblasts are characterized by the onset of PT development, which is visible via the NHS staining of the PT (**Fig. 4C**). During the sporoblast stage, the microtubules remain cytoplasmic where they form a cage-like structure that runs along the anterior-posterior axis of the parasite (**Fig. 4C**). This cytoskeletal structure may contribute to the 3-dimensional, bean-shape of the sporoblast that has previously been characterized (17).

**Figure 4.**
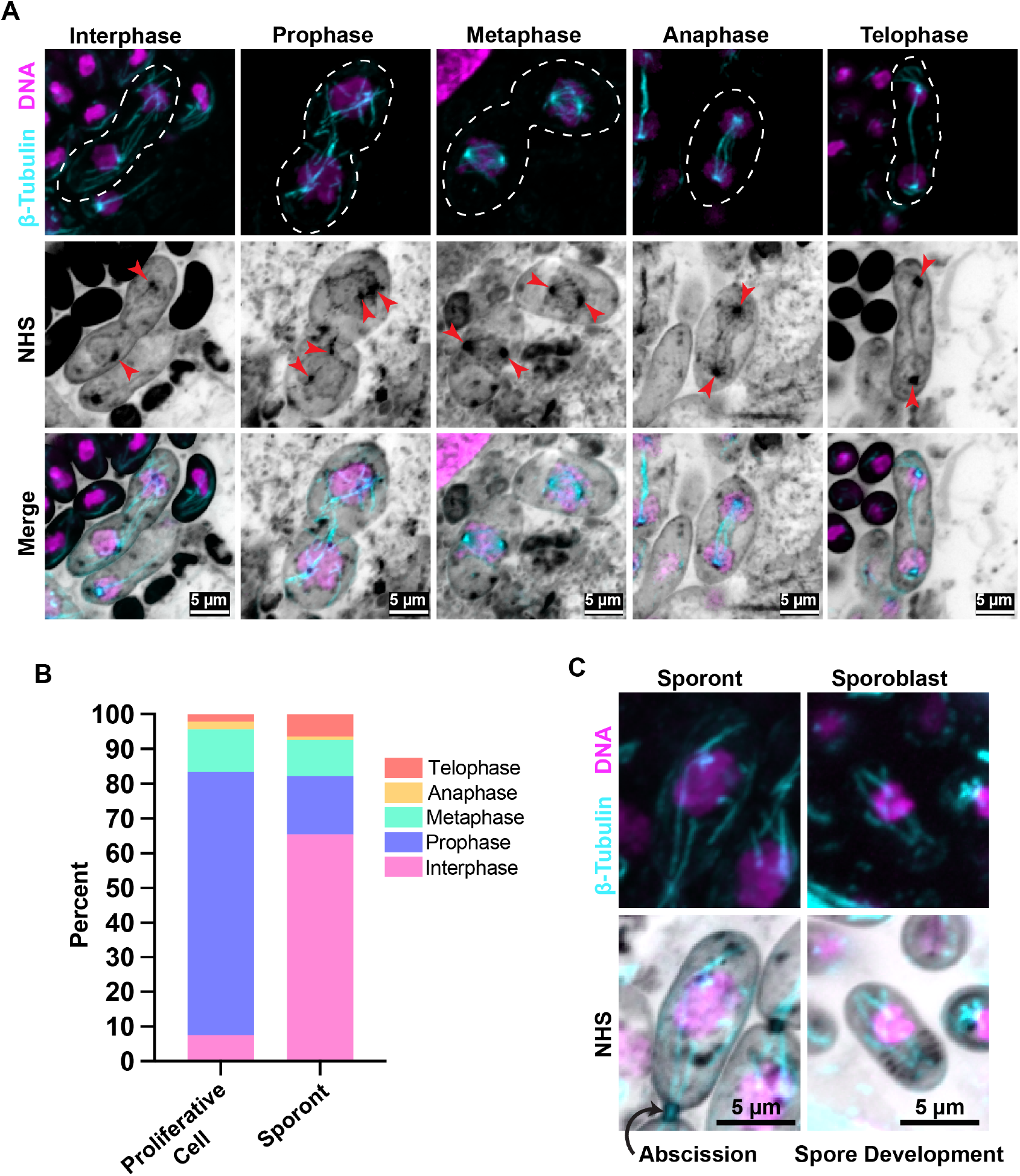
*E. intestinalis* cell cycle observed using U-ExM. A. Representative maximum intensity projections of *E. intestinalis* at each stage of the cell cycle. Cells are stained for DNA (DRAQ5, magenta), protein (NHS-ester, grayscale), and *E. intestinalis* β-tubulin (custom antibody, cyan). Red arrowheads point to spindle pole bodies. **B**. Quantification of the number of proliferative cells (*n* = 795) and sporonts (*n* = 298) at each cell cycle stage. **C**. Representative maximum intensity projections of *E. intestinalis* cytoplasmic β-tubulin in late sporonts and in sporoblasts. In both stages, β-tubulin can be observed running along the anterior-posterior axis of the parasites. All images were deconvolved in the Nikon NIS Elements AR software using the Blind deconvolution method (3 iterations).

### *E. intestinalis* polar tube development

A hallmark of sporoblast development is the appearance of the PT invasion organelle. The PT is a complex organelle composed of a linear segment at the anterior end of the spore, and a series of coils at the posterior end (**Fig. 5A, Movie S4**). How the PT assembles and develops remains an open question. Serial block-face scanning electron microscopy (SBF-SEM) data has provided a framework for PT development (17). However, due to the low throughput of SBF-SEM, only 4 infected cells were captured, and while several stages of PT development were captured, intermediate or short-lived stages were not well represented. Using U-ExM we are able to capture several stages of PT development from 296 sporoblasts across 19 cells. To track PT development, we use NHS staining along with an antibody previously identified to label the PT (anti-Eint_070340) (25). The earliest stage of PT development we capture is a short segment of the PT attached to the nucleation center (91 sporoblasts), as previously observed (17) (**Fig. 5B**). The short PT segment is typically biased towards one pole of the sporoblast, and not directly at the center of the cell. The short segment is also always in close proximity to a protein dense organelle, potentially the posterior vacuole or the Golgi-like vesicles that have been occasionally reported in the vicinity of the early PT (33) (**Fig. 5B**). Interestingly, at this stage we observe punctate staining of the protein dense organelle with the anti-Eint_070340 antibody, which is specific for the PT. One possibility is that the protein dense organelle is a reservoir for PT proteins, which are polymerized at the growing end of the PT during assembly (**Fig. 5B**). Capturing the intermediate stages between the short PT segment and PT with coils already formed has been challenging as this stage is seemingly transient. In our U-ExM dataset, we capture these early intermediate stages of the developing linear portion of the PT, prior to PT coil formation (**Fig. 5B**). These data clearly demonstrate that the linear portion of the PT grows first, followed by the assembly of the coiled region, as previously hypothesized. Interestingly, from the earliest stage of PT development, the protein dense organelle is typically localized to one pole of the sporoblast, while the nucleus is observed towards the other pole, and the nucleation center is sandwiched in between. As the PT grows, the protein dense organelle remains associated with one end of the PT at the cell pole, while the nucleation center tracks with the other end of the growing PT, ending up at the opposite end of the sporoblast. It is possible that the polarity of the cell may already be determined as early as the first appearance of the protein dense organelle. As the PT develops, the length and number of PT coils increases (**Fig. 5C**). We captured 42 sporoblasts with one PT coil, 9 sporoblasts with two PT coils, 26 sporoblasts with three PT coils, 57 sporoblasts with four PT coils, and 27 sporoblasts with five PT coils. Five PT coils is the maximum number of coils we observe. At this stage, the sporoblast likely transitions into the spore stage. The spore stage contains a thick spore wall, which is recalcitrant to expansion. Adapted protocols to allow for expansion of the spore stage (29) will likely be needed to study organelle arrangement in fully mature spores. As the spore matures and the PT coils increase, the nucleation center and the protein dense organelle at the posterior end of the spore shrink. Analyzing a large number of *E. intestinalis* parasites using U-ExM allows us to propose a generalized model for how the PT develops, in which PT proteins may first be concentrated at the protein dense organelle, before being added to the growing end of the PT. Consistent with previous data (17), the linear portion of the PT forms first, which spans the length of the sporoblast, then subsequently the coiled regions are assembled (**Fig. 5B and C)**.

**Figure 5.**
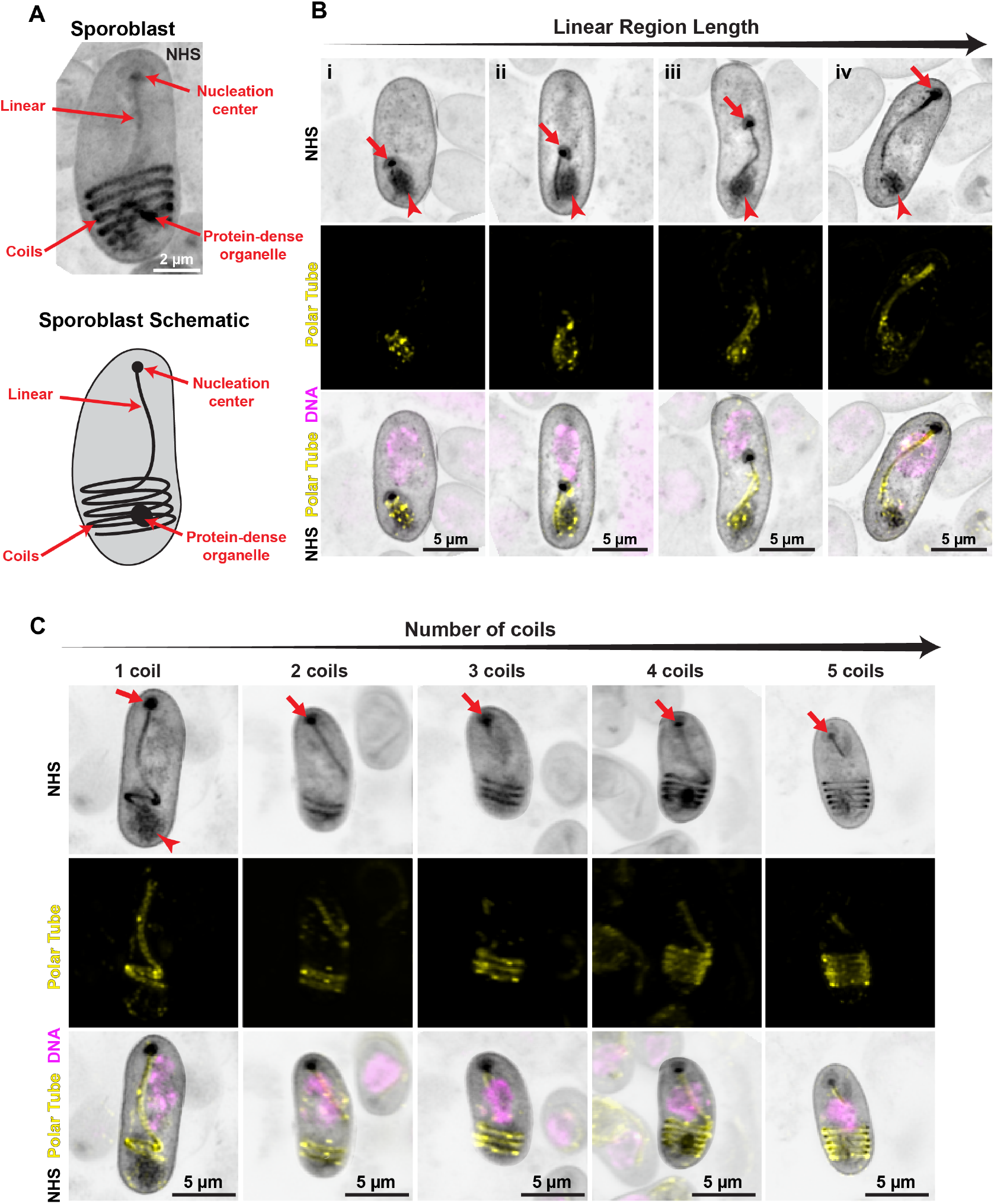
Development of the polar tube development in *E. intestinalis* visualized by U-ExM A. Representative maximum intensity projection, NHS stain only (top panel), and a corresponding schematic (bottom panel) depicting the organization of the PT in a single *E. intestinalis* parasite visualized via U-ExM. **B**. Development of the linear region of the PT across *E. intestinalis* sporoblasts, divided into four sequential states (i-iv). Representative maximum intensity projections of individual parasites with different lengths of the linear portion of the polar tube visible. Infected Vero cells were stained for DNA (DRAQ5, magenta), protein (NHS-ester, grayscale), and a previously identified protein that localizes to the polar tube (25), Eint_070340 (custom antibody, yellow). Arrows point to the nucleation center and arrowheads point to the protein dense organelle. Different lengths of the growing linear portion of the polar tube enable staging of its development. 135 sporoblasts were analyzed and assigned to the states shown in panel B (i-iv), *n*_*i*_ *=* 91; *n*_*ii*_ *=* 9; *n*_*iii*_ *=* 14; *n*_*iv*_ *=* 21. All images were deconvolved in the Nikon NIS Elements AR software using the Richardson-Lucy deconvolution method (3 iterations) **C**. Development of the coiled region of the PT across *E. intestinalis* sporoblasts. Representative maximum intensity projections of individual parasites with increasing numbers of polar tube coils visible. Infected Vero cells were stained for DNA (DRAQ5, magenta), protein (NHS-ester, grayscale), and Eint_070340 (custom antibody, yellow), same datasets as (**B**). 161 sporoblasts were analyzed and assigned to the states shown in panel C, 1 coil, *n* = 42; 2 coils, *n* = 9; 3 coils, *n* = 26; 4 coils, *n* = 57; 5 coils, *n* = 27. A gray mask with 65% opacity was placed around each sporoblast in panels A and B to highlight the sporoblast of interest. All images were deconvolved in the Nikon NIS Elements AR software using the Richardson-Lucy deconvolution method (3 iterations).

### Fusion and fission of *E. intestinalis* PVs

Having gained insight into replication dynamics using silhouette microscopy, and the associated subcellular architecture of developing parasites using U-ExM, we turned to studying the PV and its association with parasites through the life cycle. The origin of the PV surrounding parasites remains to be determined, but the leading hypothesis is that upon entry into the host cell, the sporoplasm resides within the host-derived PV membrane. At the single cell stage, electron microscopy data has shown that a membrane tightly surrounds the parasite, although it is often hard to distinguish between the PV membrane and the parasite plasma membrane (15, 16). During the replicative stages, we observe *E. intestinalis* growing in chains as opposed to spacious vacuolar compartments. This suggests that the PV membrane is tightly surrounding the parasites not only at the single cell stage, but also during early replication (**Fig. 2A and F, and Fig. 6A**). Many vacuole-residing pathogens, such as *Leishmania*, also replicate within tight-fitting membranes, known as “tight PVs” (34, 35). Interestingly, as *E. intestinalis* goes through several rounds of division, the PV transforms from a tight PV into a spacious vacuolar compartment or “spacious PV” (34), wherein the tight association between the parasites and the PV membrane is lost. For both Vero and Caco-2 host cells, we initially observe fluorescent mScarlet3 signal in between the individual chain links of parasites. As the PV transitions into a spacious PV, mScarlet3 is excluded from between the parasites, causing the individual cells to no longer be visible. The PV silhouette, now encompassing all of the individual cells, becomes a single dark shape (**Fig. 6A, Fig. S5A, and Movies S5 and S6**). Our observations are consistent with those reported for other pathogens that replicate within a vacuole. *Brucella spp*., for example, enter the host cell within a tight PV, and, as they divide, the PV divides with the pathogen creating individual pathogen-containing vacuoles. The individual pathogen-containing PVs later fuse together to create large spacious vacuoles (36).

**Figure 6.**
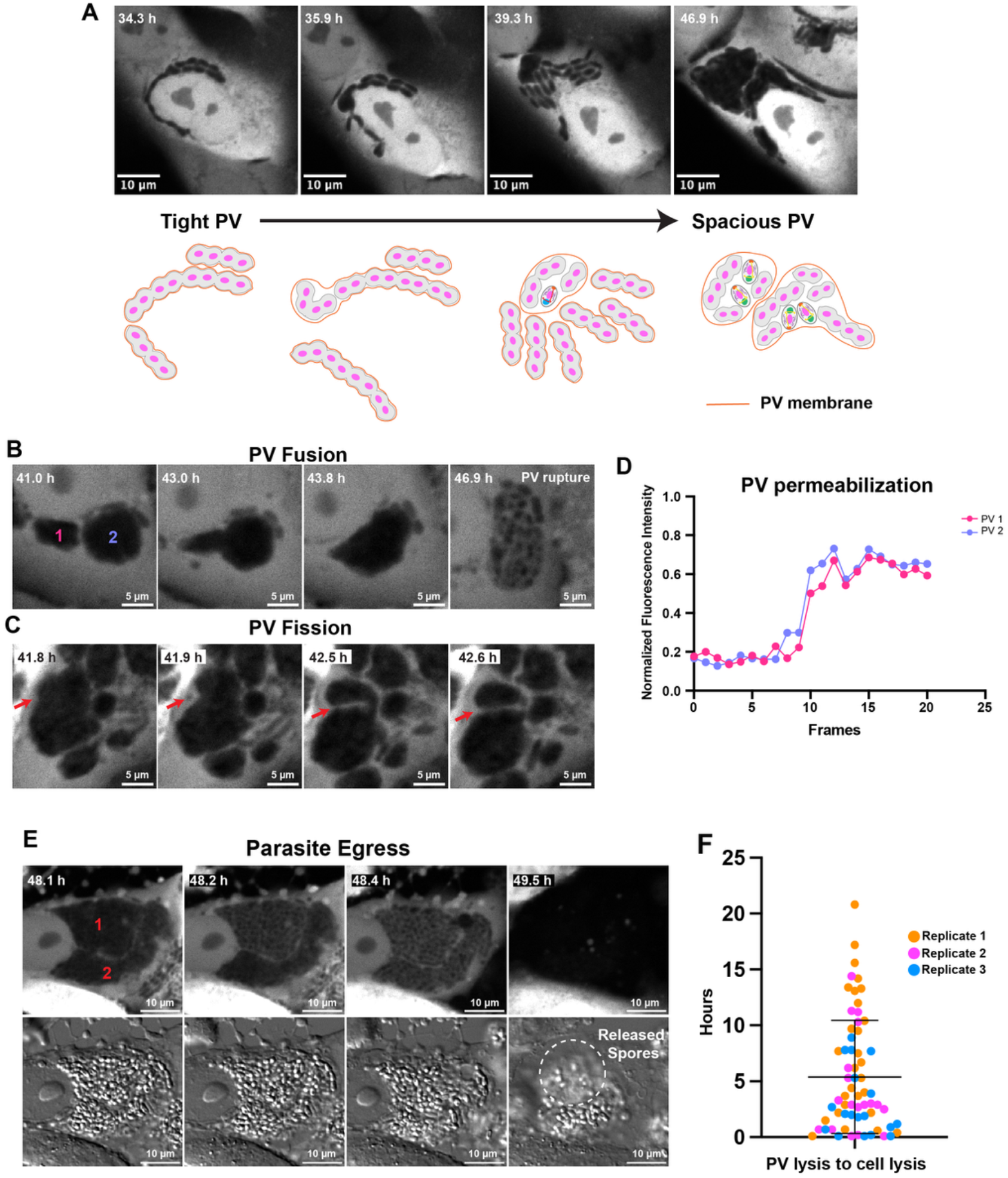
PV dynamics during *E. intestinalis* infection. A. Movie frames of *E. intestinalis* infection in Vero cells using silhouette microscopy, showing the transition from a tight PV to a spacious PV. **B, C**. Movie frames of *E. intestinalis*-infected Vero cells, showing apparent PV fusion (**B**) and fission (**C**), observed by silhouette microscopy. Numbers in (**B**) correspond to two PVs at 41.0 h that appear to fuse by ∼43.0 h. Arrow in (**C**) marks the point of fission between PVs. **D**. Representative graph showing quantification of changes in fluorescence intensity during PV permeabilization, corresponding to (**B**). Fluorescence intensity of fused PVs were measured 10 frames before and after PV permeabilization (See Materials and Methods and Supplementary Figure 6). Intensity change is very similar in regions corresponding to PV 1 and PV 2, suggesting that the two PVs have fused, and lyse as a single compartment. 25 events were analyzed with similar results. **E**. Silhouette microscopy and corresponding DIC images from movie frames of *E. intestinalis*-infected Vero cells. Timepoints are later in infection, showing PV permeabilization, followed by host cell lysis and parasite egress. All fluorescent images are representative minimum intensity projections. **F**. Quantification of the timing from PV lysis to host cell lysis. Measurements were made on cells containing 1-5 PVs per cell. The average time between PV lysis and host cell lysis is 5.4 ± 5.1 h. Three independent experiments were performed, and the error bar represents the SD across the experiments. Replicate 1, *n* = 28 PVs; Replicate 2, *n* = 18 PVs; Replicate 3, *n* = 18 PVs.

Based on observations in these other pathogens, we investigated whether the PV membrane surrounding *E. intestinalis* may also undergo fission and/or fusion. To investigate the fusogenic properties of the PV membrane, we monitored PV dynamics throughout infection. The presence of numerous vesicles that appear as silhouettes in the Caco-2 cells (**Fig. S1**), coupled with the movement of these cells during imaging, make it challenging to monitor PV dynamics. Therefore, to address the question, we analyzed PV dynamics in Vero cells only. We observed apparent fusion not only between tight PVs to create spacious PVs but also apparent fusion between multiple spacious PVs to create larger PVs as well (41 total fusion events across 220 spacious PVs in Vero cells) (**Fig. 6A and B, Movie S7, and Dataset S1**). Consistent with our observations, previous studies using transmitted light microscopy have suggested that *E. cuniculi*-containing PVs can undergo fusion (37). However, differentiating between two PVs residing close together and PVs that have truly fused is challenging, particularly using brightfield imaging alone.

While imaging PVs using silhouette microscopy, we noticed that late in infection, the fluorescence of PVs rapidly increases, likely due to rupture or leakage of the PV membrane and diffusion of cytoplasmic mScarlet3 into the PV (**Fig. 6B and Movie S7**). This phenomenon provides a serendipitous tool to differentiate between PVs in close proximity versus PVs that have fused. Conceptually, the phenomenon is similar to a fluorescence recovery after photobleaching (FRAP) experiment: The PV is dark (analogous to being bleached), and fluorescence rushes in (analogous to fluorescence recovery). If PVs have truly fused, the parasites within the PV are encompassed by a single membrane, and upon PV membrane permeabilization, mScarlet3 fluorescence should increase at the same time across the fused compartment. In contrast, if PVs are in close proximity to each other but not fused, the increase in mScarlet3 fluorescence should not be correlated, and the fluorescence should increase at different times in each PV (**Fig. S6A and B**). We analyzed the fluorescence intensities of PVs that resulted from apparent fusion events and then underwent membrane permeabilization (25 fusion events in Vero cells) (see Materials and Methods). Our analysis reveals that PVs that have fused, have almost identical kinetics of “fluorescence recovery” suggesting that the PVs are truly fusing, and are not simply in close proximity (**Fig. 6D and Fig. S6C**). While fusion between spacious PVs occurs with some frequency, most spacious PVs in a cell do not fuse with one another (178 spacious PVs do not fuse across 220 PVs). It is unclear what renders the PVs fusion-competent, but one possibility is that there may be temporal regulation of the PV membrane composition. If the lipid composition of the PV membranes change over time, the PV membrane may acquire different parasite or host markers, which may modulate their fusogenic properties. In addition to fusion, fission of spacious vacuoles also occurs but at a lower frequency (9 fission events across 220 spacious PVs in Vero cells) (**Fig. 6C, Movie S8, and Dataset S1**). However, fission of tight vacuoles is also likely occurring while the parasites are within tight vacuoles. During abscission, we observe the chains of parasites breaking apart from one another. If there is a PV membrane surrounding the chains of parasites, each time the parasites undergo abscission and separate from the rest of the chain, the PV membrane must also be undergoing fission, suggesting fission may be constantly occurring. Overall our data suggest that the PV membrane is highly dynamic during infection, and that fission and fusion play a role in PV remodeling and maintenance.

### *E. intestinalis* ruptures the PV membrane and exits via host cell lysis

Following spacious vacuole formation, *E. intestinalis* forms mature spores that are ready to exit host cells. Much of the data surrounding parasite egress comes from microsporidia infection in nematodes (38, 39). Our current understanding of how human infecting species, including *E. intestinalis*, exit the host cell remains poorly understood. Using silhouette microscopy we monitored parasite egress in Vero and Caco-2 cells in real time. In the late stages of the life cycle, the PVs are full of mature spores ready for exit. Approximately 43 hpi we observe mScarlet3 signal rushing into the PV compartment and surrounding individual parasites (**Fig. 6E**). This suggests that the PV membrane is ruptured and the parasites are released into the host cytoplasm. Anywhere between minutes to several hours after PV membrane rupture, the host cell lyses and mature spores are released into the environment, indicative of lytic egress (**Fig. 6E and F, Movie S9, and Dataset 1**). We observed this exit strategy in both Vero cells and Caco-2 cells suggesting this is a conserved mechanism and not limited to specific cell types (**Fig. S5B and C, Movie S10, and Dataset 2**). In other PV residing pathogens, such as *Toxoplasma gondii* and *P. falciparum*, pore-forming proteins and proteases are secreted to initiate PV membrane rupture and host cell lysis (40–42). It is possible that similarly, upon maturation, *E. intestinalis* may secrete proteins to facilitate PV membrane rupture and host cell lysis. Interestingly, scRNA-seq data has revealed that late in infection, expression of secreted proteins is increased, followed by host cell death via pyroptosis (25). Combined with our live-cell imaging data, this suggests that as parasites mature, they may secrete proteins in order to rupture the PV membrane and the host cell plasma membrane in order to facilitate egress. This exit strategy is in contrast to the previously described exit strategy employed by another microsporidian species, *N. parisii. N. parisii* uses a vesicle-mediated, non-lytic mechanism to exit intestinal cells in *C. elegans* (43). In contrast to *E. intestinalis, N. parisii* does not replicate within a PV but rather replicates directly in the host cytoplasm. It is possible that *E. intestinalis* may use a different exit strategy *in vivo* than what we observe in our 2D cell culture models. Alternatively, it is possible that microsporidian species replicating in different niches may use different mechanisms to exit the host cell.

## Discussion

Together, silhouette microscopy and U-ExM synergize to paint a dynamic picture of the *E. intestinalis* life cycle, from entry to exit, with specificity and throughput that resolves the parasite cytoskeleton and polar tube development (**Fig. 7**). Below, we discuss some aspects of the *E. intestinalis* life cycle and our imaging tools.

**Figure 7.**
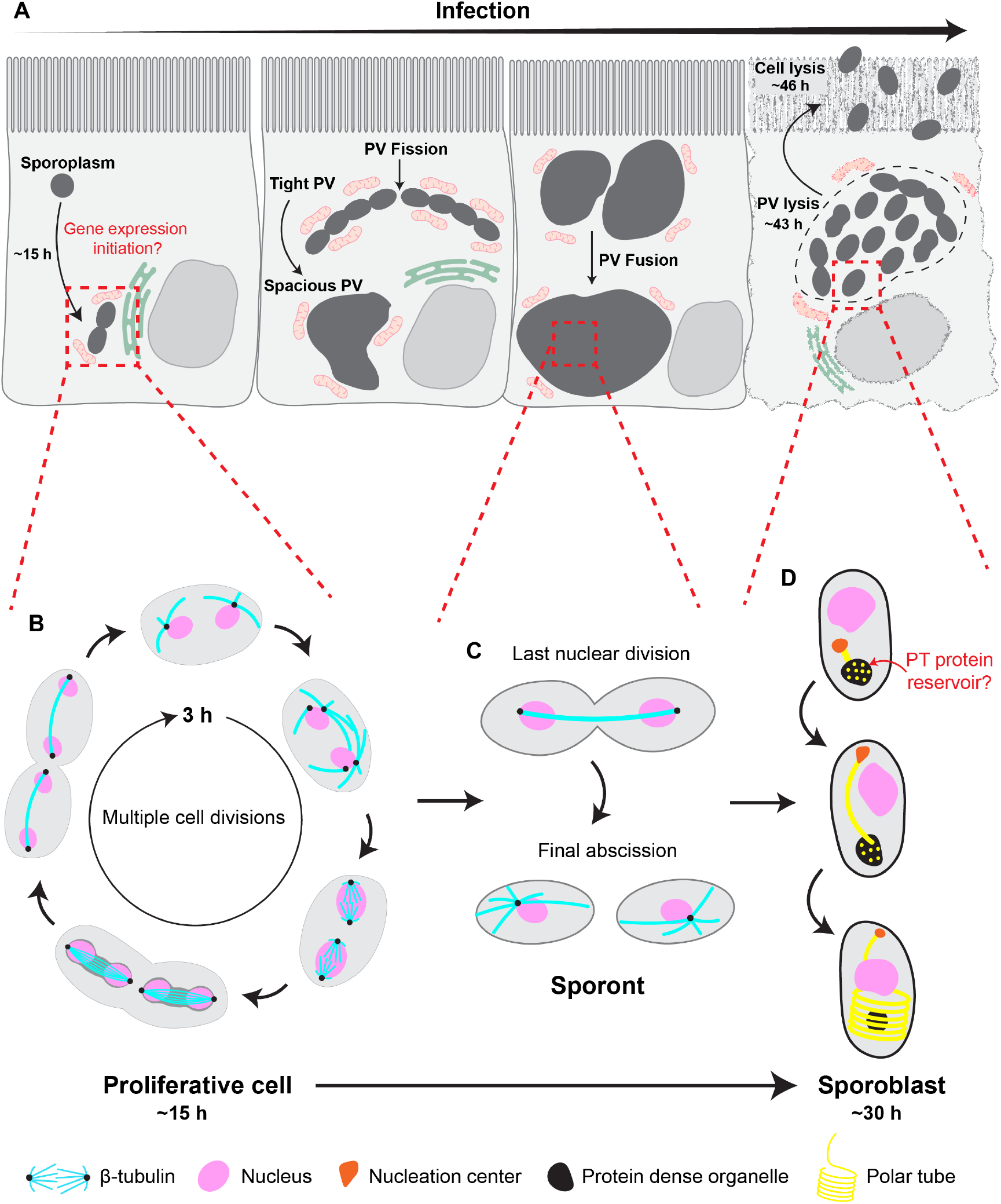
Dynamics of the *E. intestinalis* intracellular life cycle. A-D. Schematic overview of the intracellular life cycle of *E. intestinalis*. **A**. Upon entry of the sporoplasm into the host cell, *E. intestinalis* undergoes a lag phase of ∼15 h before completing the first division. Following the lag phase, *E. intestinalis* replicates within a tight PV which is constantly undergoing fission. The tight PVs may enable efficient nutrient acquisition from host organelles during rapid replication. As the nutrient requirements of the parasites decrease, the tight PVs transition into spacious PVs. Spacious PVs can undergo fusion to create larger vacuoles. Approximately 43 hpi, the PVs rupture and parasites are released into the host cytoplasm. Approximately 46 hpi, the host cell lyses, facilitating parasite egress. **B**. Proliferative cells undergo several rounds of cell division and are responsible for most parasite replication. **C**. Upon entering the sporont stage, the spore wall begins to form. The parasites undergo one last cell division and complete the final abscission. **D**. The sporoblast stage begins upon PT development. PT proteins concentrate at the protein dense organelle and are added onto the growing end of the PT. The linear portion of the PT grows first followed by the coiled region.

### *E. intestinalis* lag time

Upon entry of the sporoplasm, *E. intestinalis* undergoes a lag time of ∼14-15 h before replication begins (**Fig. 2, Fig. 7A**). Undergoing a lag time post invasion is common among intracellular pathogens (44, 45) and may be due to the time necessary for initiating gene expression, adapting to the host cell environment, or hijacking host machinery for replication. *Salmonella typhimurium*, for example, undergoes a lag time of 4 h during which the bacteria are recruiting host factors, such as lysosomal membrane glycoproteins, to the PV (46). Only upon recruitment of host factors can the bacterial cells begin to replicate. Due to the small genome size of *E. intestinalis*, these parasites lack the ability to synthesize nucleotides, amino acids, and lipids, requiring a host for metabolites. As a result, the lag time post invasion may be due to recruiting host factors and/or metabolites to the PV in order to initiate gene expression and facilitate growth, similar to *S. typhimurium*. Previous studies have found that *E. intestinalis* transcript levels are low after invasion and increase substantially by 12 hpi. The genes initially expressed are housekeeping genes, transcription initiation factors, and protein synthesis machinery (25). This suggests that *E. intestinalis* requires several hours to initiate gene expression in preparation for replication. Alternatively, the lag phase may be a consequence of the microsporidian invasion mechanism, which is rapid. On the timescale of just a few hundred milliseconds, the PT extends rapidly, and the sporoplasm, including the nucleus, is massively deformed and transported at high velocity through the PT, into the host cell (11). This deformation may cause damage to the nucleus and other sporoplasm cargo and require repair upon entry into the host cell. Membrane repair can be on the order of seconds (47) to hours (48) depending on the extent of the damage, but it is quite likely to be substantial, given the mode of cargo transport (49). Nuclear deformation is also known to lead to DNA damage and chromosome breakage (50, 51), which may require repair before DNA replication can begin. Therefore, the lag time upon invasion may be a result of sporoplasm repair following invasion, or any combination of sporoplasm repair, hijacking host machinery, and initiation of gene expression to facilitate replication.

### *E. intestinalis* PV dynamics

The PV is essential for protecting the parasites from the host cytoplasm, but it also presents a barrier for nutrient acquisition from the host. Here we show that the PV has dynamic properties and can undergo fusion and fission under our experimental conditions (**Fig. 6, Fig. 7A**). During proliferation, the PV remains tightly associated with the parasite plasma membrane and during each parasite division, the PV membrane undergoes fission with the parasites. This suggests that the PV membrane is undergoing fission constantly during replication, a time in which rapid growth and expansion is occurring. PV fission during the proliferative phase may be advantageous to the parasites, in that fission results in an increased surface area to volume ratio, facilitating increased accessibility for interactions with host organelles to acquire nutrients. As tight PVs fuse to form spacious PVs, the surface area to volume ratio decreases. This likely reflects the reduced nutrient requirements of the parasites near the end of their life cycle. Similar vacuole fusion has also been observed in other pathogens such as *Chlamydia trachomatis* (52). In this case, fusion has been hypothesized to be important for nutrient sharing and/or genetic exchange, and it has been shown to be important for virulence. Sex-related genes have been identified in *Encephalitozoon* species which suggests genetic exchange may occur between microsporidia (53), perhaps during mixed infections, which may also be facilitated by PV fusion. The lipid composition of *C. trachomatis* inclusions (PVs) was found to play a significant role in fusion (54). Depleting the lipid PI(3,4)P_2_ or sphingomyelin causes delays in fusion. Interestingly, sphingomyelin maintains the organization of lipid rafts which are required for membrane fusion. Currently little is known about the mechanism of PV fusion for microsporidia. Monitoring the lipid composition of *E. intestinalis* PVs, how this changes over time, and how perturbing the fusogenic PV properties impacts infection has the potential to provide insight into the changes in fusion-competence we observe, and a better understanding of how fission and fusion may be related to virulence.

### Expanding the imaging toolkit to study microsporidia

Here we developed silhouette microscopy which establishes a method for sustained, live-cell imaging in microsporidia, overcoming the challenge posed by the lack of genetic tools, which precludes fluorescent tags for live-cell imaging. Silhouette microscopy enables the visualization of parasite morphology during replication at the single parasite level, allowing us to define the kinetics of parasite division, as well as characterize PV dynamics, and parasite egress. The simplicity of the technique makes it an excellent platform to study life cycle dynamics and kinetics for other microsporidian species in the future. In the context of drug treatments or other perturbations to the host cell, silhouette microscopy can be used to monitor parasite morphology and changes in life cycle kinetics. This line of experiments would provide insights into the stages of the parasitic life cycle that may be modulated by different drug treatments. In addition, silhouette microscopy can be coupled with live host cell markers to label host organelles and proteins of interest to visualize and study host-parasite interactions.

Microsporidian cell biology is challenging to study using optical microscopy, due to its small size. While many EM studies have provided invaluable insight into cellular level ultrastructure (15, 17), the throughput of EM is low, often impeding rigorous statistical analyses. Moreover, labeling specific proteins for EM is arduous, often resulting in ambiguity of the protein identity. To expand our imaging toolkit, and bridge the resolution gap between conventional immunofluorescence and transmission electron microscopy imaging, we adapted a previously established U-ExM protocol to study the organization and development of *E. intestinalis* cytoskeleton and organelles in infected cells. Our workflow enables high throughput (1622 parasites analyzed in this work) with the specificity resulting from antibody staining, to provide insights into cytoskeletal organization and organelle development (**Fig. 7B-D**). Using the U-ExM workflow will be valuable for studying the localization of microsporidian proteins at the ultrastructural level, and using this information to hypothesize functions of unstudied proteins.

## Materials and methods

### Maintenance of mammalian cell lines

Vero cells (ATCC CCL-81), Caco-2 cells (Millipore sigma 86010202), and HEK293FT cells (gift from Susan Smith, Skirball Institute) were grown at 37 °C in a humidified atmosphere with 5% CO_2_ in DMEM high glucose growth medium (DMEM-HG) (ThermoFisher Scientific 11965092) supplemented with 10% heat inactivated fetal bovine serum (FBS) (VWR Life Science 89510-188) and non-essential amino acids (NEAA) (Gibco 11140050). All host cell and parasite lines are mycoplasma-negative, and were tested monthly using the Minerva Biolabs Venor GeM classic mycoplasma detection kit (11-1100G).

### Propagation of *E. intestinalis* spores

*E. intestinalis* spores (ATCC 50506) were propagated in Vero cells. Vero cells were grown in a 75 cm^2^ tissue culture flask using DMEM-HG supplemented with 10% FBS at 37 °C and with 5% CO_2_. At 70%-80% confluence, the media was switched to DMEM-HG supplemented with 3% FBS and NEAA, and *E. intestinalis* spores were added. Infected cells were allowed to grow for 8–12 days, until the Vero cells began to lyse, and the medium was changed every two days. Spores were purified as follows, using a previously established protocol (55). To purify spores, the infected cells were detached from tissue culture flasks using a cell scraper and moved to a 50 mL conical tube, followed by centrifugation at 2000 × g for 10 min at 25 °C. Cells were resuspended in 12 mL 1X DPBS (Thermo Fisher 14190250) + 0.1% SDS and incubated on ice for 10 min with vortexing every 5 min. The suspension was divided into two 15 mL conical tubes. The released spores were purified using a 2-step Percoll gradient (55). Equal volumes (6 mL) of spore suspension and 100% Percoll were added to a 15 mL conical tube, vortexed, and then centrifuged at 4000 × g for 30 min at 25 °C. The spore pellets were washed 3 times with 1X DPBS. The spores were further purified via Percoll gradient (25%, 50%, 75%, 100%) centrifugation. To do this, spores were layered over the Percoll gradient and centrifuged at 7000 xg for 30 min at 25 °C (Bioflex HC rotor, ThermoFisher Scientific). Bands containing various impurities remained in the supernatant/gradient fractions, and were removed using a vacuum. The pellet at the bottom of the tube, containing mature spores, was washed with 1X DPBS and ultracentrifuged again at 7000 xg for 30 min at 25 °C. The spore pellet was resuspended in 1X DPBS and transferred to a 1.5 mL microcentrifuge tube and centrifuged at 4000 × g for 5 min at 25 °C. The pellet was washed 2 additional times with 1X DPBS. The final pellet was resuspended in 1X DPBS and stored at 4 °C.

### Generation of mScarlet3 cell lines

Vero-mScarlet3 and Caco-2-mScarlet3 cells were generated by lentiviral transduction. Lentivirus was produced in HEK 293FT cells co-transfected with 5.8 µg psPAX2 (Addgene #12260), 2.9 µg pMD2.G (Addgene #12259) (plasmids psPAX2 and pMD2.G were gifts from Didier Trono) and 5.8 µg pBEL2982 (derivative of pLV-mitoDsRed (gift from Pantelis Tsoulfas; Addgene #44386) containing lentiviral backbone expressing mScarlet3; Addgene #263107) using Lipofectamine 2000 (Thermo Fisher 11668027) according to the manufacturer’s instructions. 48 h after transfection, the virus-containing supernatant was collected and centrifuged at 500 xg for 5 min at room temperature to pellet cell debris. The supernatant was then filtered with a 0.45 µm filter, supplemented with 1 µg/ml of polybrene (Millipore sigma TR-1003-G) and added to target cells. ∼24 h after transduction, the virus was removed and replaced with fresh DMEM-HG supplemented with 10% FBS and NEAA.

### Live-cell imaging and analysis

Vero and Caco-2 cells stably expressing mScarlet3 were cultured in a 35 mm glass bottom imaging dish (Ibidi 50-305-807) or a 6 well glass bottom imaging dish (Cellvis P06-1.5H-N) in DMEM-HG supplemented with 10% FBS and NEAA. Immediately before imaging, the media was replaced with DMEM-HG supplemented with 3% FBS and NEAA, and infected with *E. intestinalis* spores at an MOI of 50. Live imaging was performed on a Nikon CSU-W1 SoRa spinning disk confocal microscope equipped with Nikon Perfect Focus. Cells were maintained at 37 °C and 5% CO_2_ with a stage top incubator with flow control and a sub stage environmental enclosure (Tokai hit). Cells were imaged by DIC and fluorescence microscopy (577 laser) at 60X using a Nikon Apo 60x 1.42 oil objective or Apo 20x 0.8 objective with 2.8x magnification. Z-stacks were acquired with a 0.2 µm or 0.3 µm step size every 7 minutes at 10 different fields of view. The fields of view were selected at random, as host cells were not visibly infected at the start of the imaging experiment. To analyze the parasite life cycle, later frames of the movies were examined to identify clearly infected cells. Using these frames as a starting point, the movie was tracked backwards to capture the entry of a single sporoplasm, which appears as a single silhouette. Parasites were only quantified if entry of the sporoplasm could be identified. Three experiments were performed. Minimum intensity projection images and movies were created in the Nikon NIS Elements AR software. All images and movies were cropped in Fiji (56).

PV fusion “FRAP” analysis was performed in Fiji (56). Briefly, a 20×20 pixel region of interest (ROI) for each PV that underwent fusion was selected using the oval selection tool. The fluorescence intensity for each PV was measured 10 frames prior to PV permeabilization and 10 frames after PV permeabilization. A 20×20 pixel ROI of the host cell cytoplasm (outside of the PV) was measured as the “cytoplasm” fluorescence intensity and a 20×20 pixel ROI outside of the cell was measured as the background fluorescence. The cytoplasmic and background fluorescence intensities were measured for each cell. The PV fluorescence intensities were normalized to the cytoplasmic fluorescence intensities and background fluorescence intensities and plotted using GraphPad Prism version 11.0.0.

### RNA FISH and calcofluor white staining

RNA FISH and calcofluor white staining in conjunction with DIC imaging was used to assess approximately when parasites begin to mature. The RNA FISH probe labels *E. intestinalis* 16S rRNA of actively replicating parasites and calcofluor white labels chitin of maturing parasites. Using fluorescence and DIC, we can correlate the parasite stages. Vero cells were seeded onto coverslips and infected at an MOI of 30 for 30 h. The cells were washed once in 1X PBS (37 mM NaCl, 2.7 mM KCl, 10 mM Na_2_HPO_4_, 1.8 mM KH_2_PO_4_, pH 7.4) and fixed in 4% PFA in PBS at room temperature for 30 min. Coverslips were then washed in PBS + 0.1% Triton X-100 3 times for 10 min at room temperature and once with hybridization buffer (900 mM NaCl, 20 mM Tris HCl, 0.01% SDS). Then, 50 μL FISH staining solution (125 nM FISH probe in hybridization buffer) was added per coverslip and incubated for 18 h at 37 °C. An *E. intestinalis* 16S rRNA-specific FISH probe conjugated to Quasar 590 (LGC Biosearch Technologies) was used. Following incubation, coverslips were washed twice with wash buffer (hybridization buffer + 5 mM EDTA) for 30 min at 37 °C to remove excess FISH probe. Then, 2 µg/µL calcofluor white (18909-100ML-F) and 5 µM DRAQ5 (Novus biologicals NBP2-81125-200ul) in 1X PBS + 0.1% Triton X-100 was added, and samples were incubated for 30 min at room temperature. Coverslips were washed once with 1X PBS + 0.1% Triton X-100 and mounted onto slides with Prolong Diamond antifade (ThermoFisher Scientific; P36965) and sealed. Samples were imaged on a Nikon CSU-W1 SoRa spinning disk confocal microscope with a Nikon Apo 60×1.42 Oil objective. Z-stacks were acquired with 0.3 μm spacing. Maximum intensity projection images were created in the Nikon NIS Elements AR software. Three experiments were performed and ≥100 parasites were analyzed per experiment.

### Custom antibody generation

Custom polyclonal β-tubulin and Eint_070340 antibodies were raised at Capra Science (Sweden). The β-tubulin antibody was generated by immunizing rabbits with the peptide sequence of CDQSGRYVGTSDNQLER and the Eint_070340 antibody was generated by immunizing chickens with the peptide sequence of CAKTPESKEGAKGKEK. Polyclonal antibodies were affinity purified from antiserum or egg yolks, respectively, using columns coupled with the antigen and extraction of IgG using protein A affinity chromatography.

### Immunofluorescence microscopy of unexpanded samples

Vero cells were seeded onto coverslips and infected with *E. intestinalis* spores at an MOI of 30 for 26 h. The cells were fixed in 4% PFA in 1X PBS at room temperature for 30 min, followed by washing 3 times with 1X PBS. The cells were permeabilized and blocked in Blocking buffer (3% bovine serum albumin (BSA) in 1X PBS + 0.25% Triton x-100) for 30 min at room temperature. The cells were stained with primary antibody against *E. intestinalis* β-tubulin at 1.5 µg/ml (Capra science) in Blocking buffer overnight at 4°C. The cells were washed 3 times with 1X PBS + 0.1% Triton x-100 for 5 min each at room temperature and incubated with goat anti-rabbit Alexa 568 at 4 µg/mL (Invitrogen A-11036), DRAQ5 at 5 µM, and NHS-405 at 20 µg/mL (Thermo fisher A30000) in Blocking buffer for 30 min at room temperature. The cells were washed 3 times with 1X PBS + 0.1% Triton x-100 for 10 min each at room temperature and mounted onto slides with Prolong Diamond antifade. The slides were visualized using a Nikon CSU-W1 SoRa spinning disk confocal microscope. All images were acquired using a Nikon Apo 60x 1.42 oil objective. Z-stacks were acquired with 0.2 μm spacing. Maximum intensity projection images were created in the Nikon NIS Elements AR software. Three experiments were performed and ≥100 parasites were analyzed from each experiment.

### Ultrastructure expansion microscopy

U-ExM was performed as follows, based on previously described protocols for *Chlamydomonas reinhardtii* and *Plasmodium falciparum* (20, 21). Vero cells were seeded onto uncoated coverslips and infected with *E. intestinalis* spores at an MOI of 30, for 20, 26, or 30 h. The cells were fixed in 4% PFA in 1X PBS for 30 min at room temperature followed by washing 3 times with 1X PBS. Following fixation, the coverslips were incubated in anchoring solution (1.4% (v/v) formaldehyde (Sigma F8775-25mL)/2% (v/v) acrylamide (Sigma A4058) in PBS) overnight at 37 °C. The coverslips were washed 3 times with PBS. Monomer solution (19% (wt/wt) sodium acrylate (Sigma 408220), 10% (v/v) acrylamide, 0.1% N,N’-methylenbisacrylamide (Sigma M1533), 0.5% (v/v) tetraethylenediamine (ThermoFisher 17919), and 0.5% (w/v) ammonium persulfate (ThermoFisher 17874)) was used for gelation and the gels polymerized for 30 min at 37 °C. The gels were transferred to denaturation buffer (200 mM SDS, 200 mM NaCl, 50 mM Tris in water, pH 9) for 15 min at room temperature and then for 90 min at 95 °C. The denatured gels were expanded in MilliQ water 3 times for 30 min each at room temperature. The gels were cut into smaller pieces and washed with 1X PBS twice for 15 min at room temperature to shrink the gels. The shrunken gels were blocked with 3% BSA in 1X PBS for 30 min at room temperature. Following blocking, the gels were incubated with primary antibody against *E. intestinalis* β-tubulin at 3.7 µg/ml (Capra science) and Eint_070340 at 3.4 µg/mL (Capra science) in 3% BSA in 1X PBS overnight at room temperature. The gels were washed 3 times with 1X PBS + 0.5% Tween-20 (PBS-T) for 10 min each at room temperature. The gels were incubated with goat anti-rabbit Alexa 568 at 4 µg/mL, goat anti-chicken Alexa 488 (Invitrogen A-11039) at 4 µg/mL, DRAQ5 at 25 µM, and NHS-405 at 50 µg/mL for 2.5 h at room temperature. The gels were washed 3 times with PBS-T for 10 min each at room temperature. The gels were re-expanded in MilliQ water 3 times for 30 min each at room temperature. The gels were imaged in 35 mm glass bottom imaging dishes (Ibidi) that have been precoated with poly-D-lysine. All images were acquired with a Nikon CSU-W1 SoRa spinning disk confocal microscope using a Nikon Apo 60x 1.42 oil objective or Zeiss LSM800 AxioObserver with an Airyscan detector using a Zeiss Apo 40x 1.2 water objective. Z-stacks were acquired with a 0.18 µm -0.2 µm step size. All images taken on the Nikon spinning disk underwent deconvolution in the Nikon NIS Elements AR software using the Blind or Richardson-Lucy deconvolution methods (3 iterations) (specified in figure legends). All images taken on the Zeiss LSM800 were Airyscan processed using 3D processing with auto filtering on ZEN Blue. All images were cropped in Fiji (56). Three experiments were performed for each time point.

## Acknowledgements

We thank Noelle Antao, Atty Chang, Olivia Choi, Monica Perumattam, Paris Watson, Pengzheng Yong, and Snow Zhou for feedback on our manuscript and all members of the Bhabha+Ekiert labs for helpful discussions. We thank the NYU Microscopy Laboratory and the Integrated Imaging Center at JHU for providing expertise and access to their microscopes. We gratefully acknowledge the following funding sources: SSP-2018-2737 (Searle Scholars Program, to G.B.); R01AI147131 (National Institute of Allergy and Infectious Disease, to G.B.); Irma T. Hirschl Career Scientist Award (to G.B.); NIH Institutional training grant T32GM136542 (Training Program in Cell Biology, to K.L.M), American Heart Association Grant no. 24PRE1187590 (to K.L.M), 1F31AI183707 (National Institute of Allergy and Infectious Disease, to K.L.M), G.B. is a Pew Scholar in the Biomedical Sciences, supported by The Pew Charitable Trusts (PEW-00033055).

## Author contributions

K.L.M., G.B., and D.C.E. were responsible for project conceptualization and experimental design. K.L.M. and S.A. were responsible for data collection. K.L.M., G.B., and D.C.E. were responsible for data analysis. K.L.M, G.B., and D.C.E. were responsible for manuscript writing. G.B. and D.C.E. supervised the project. All authors provided feedback on the manuscript.

## AI statement

No AI tools were used in the preparation, analysis, or writing of this manuscript.

## Data availability

Microscopy data will be deposited in BioImage Archive and is available from the authors upon request.

