## Supplementary information for "Live-cell imaging and ultrastructure expansion microscopy reveal the dynamic intracellular life cycle of the microsporidian parasite *Encephalitozoon intestinalis*"

### Supplementary data

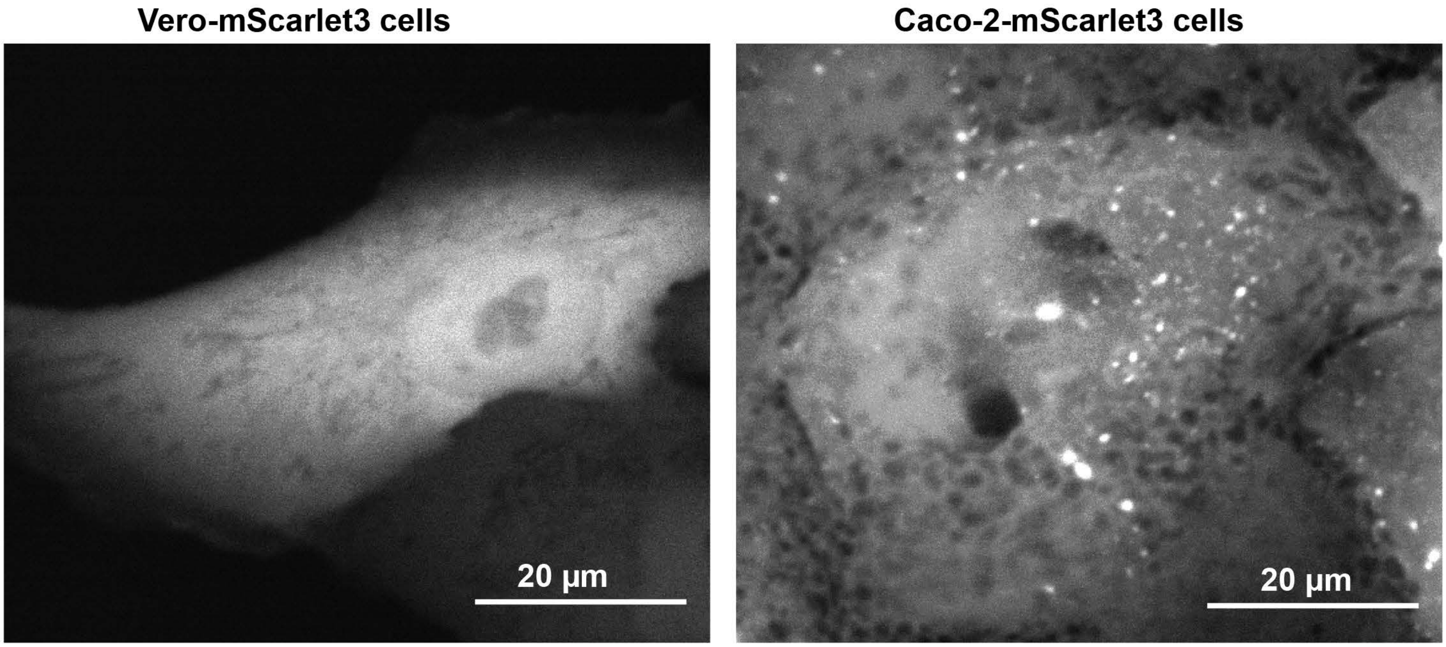


**Figure S1. Vero and Caco-2 cells transfected with mScarlet.** Representative images of Vero-mScarlet3 cells and Caco-2-mScarlet cells. Vero-mScarlet3 cells typically grow in monolayers and are relatively featureless, whereas Caco-2-mScarlet cells typically grow in clusters and commonly display dark vesicle-like structures in the cytoplasm. These features make Vero cells easier to use for silhouette microscopy analysis, while Caco-2 cells are more challenging to analyze.

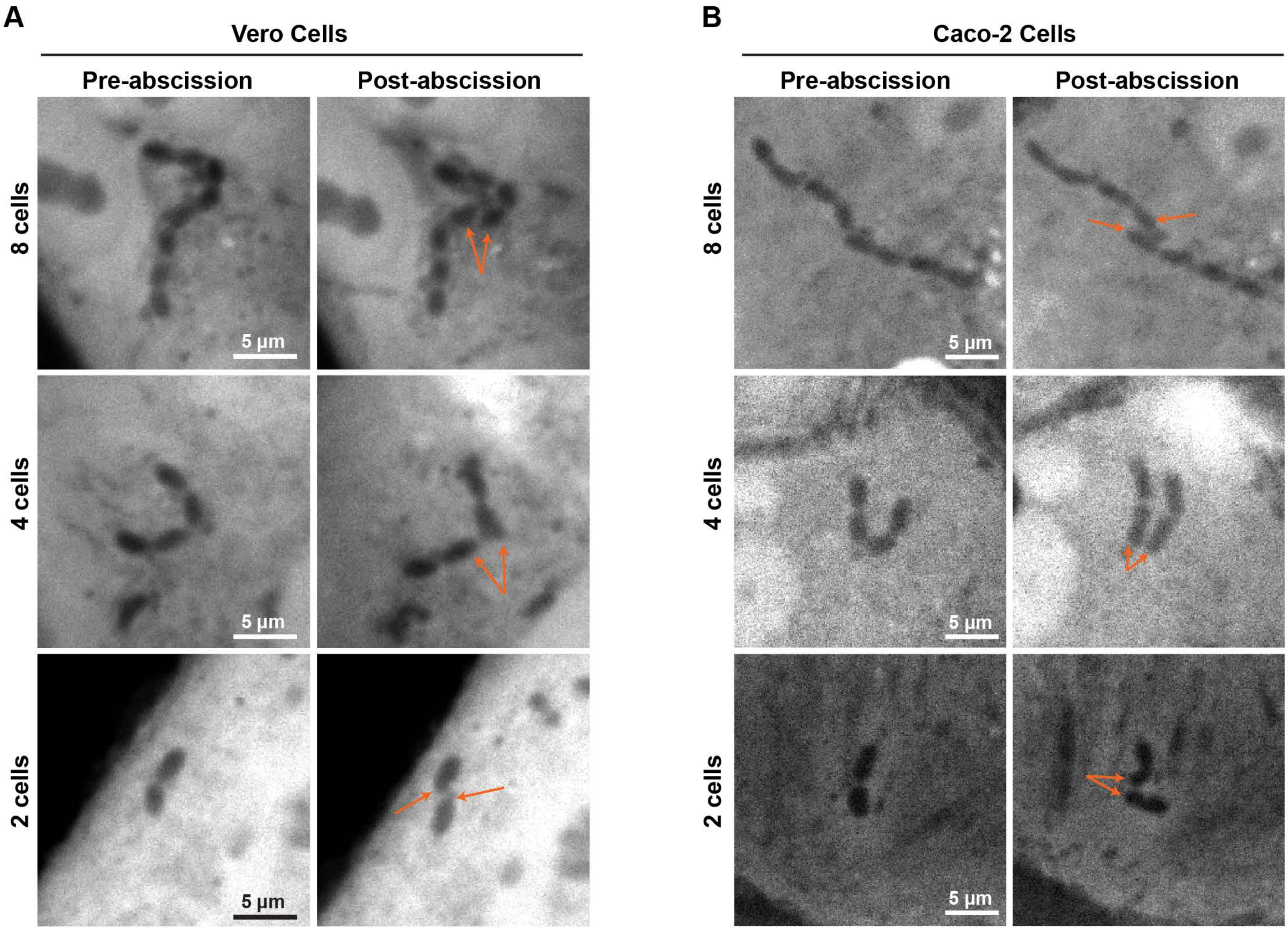


**Figure S2.** ***E intestinalis*** **chains undergoing abscission. A.** Representative images of *E. intestinalis* chains undergoing abscission at the 2, 4, and 8 cell stages in Vero-mScarlet3 cells. **B.** Representative images of *E. intestinalis* chains undergoing abscission at the 2, 4, and 8 cell stages in Caco-2-mScarlet cells

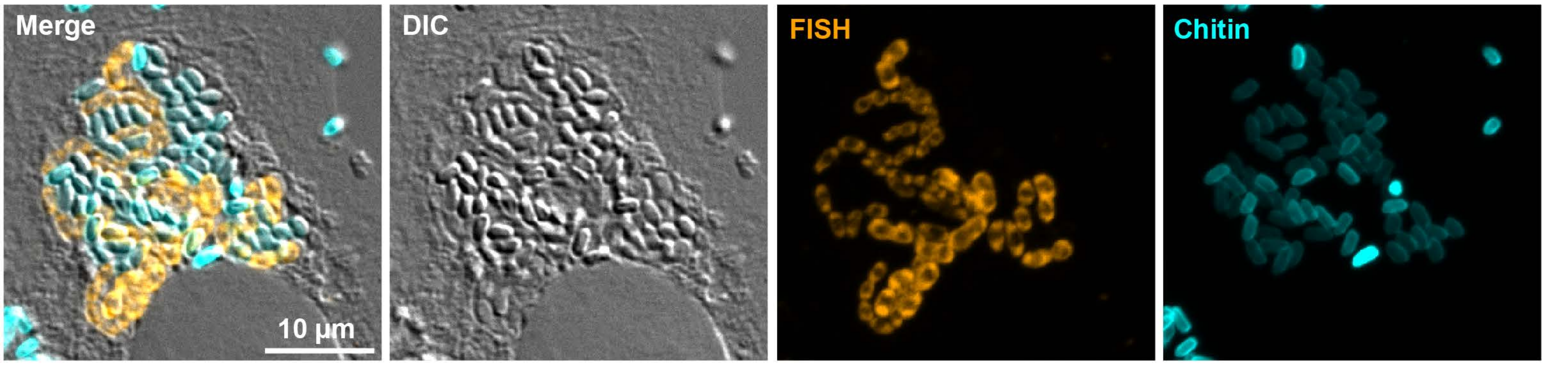


**Figure S3. Correlation of parasite contrast in DIC with chitin stain and rRNA FISH, to identify maturing spores by DIC.** Representative images of Vero cells infected with E. intestinalis for 30 h and stained for *E. intestinalis* 16s rRNA (FISH, orange) and chitin (calcofluor white, cyan), also imaged in DIC. Parasites that appear more clearly “3D” in DIC correspond to chitin-positive, FISH-negative parasites. Three biological replicates were performed. n ≥ 100 parasites were analyzed per experiment.

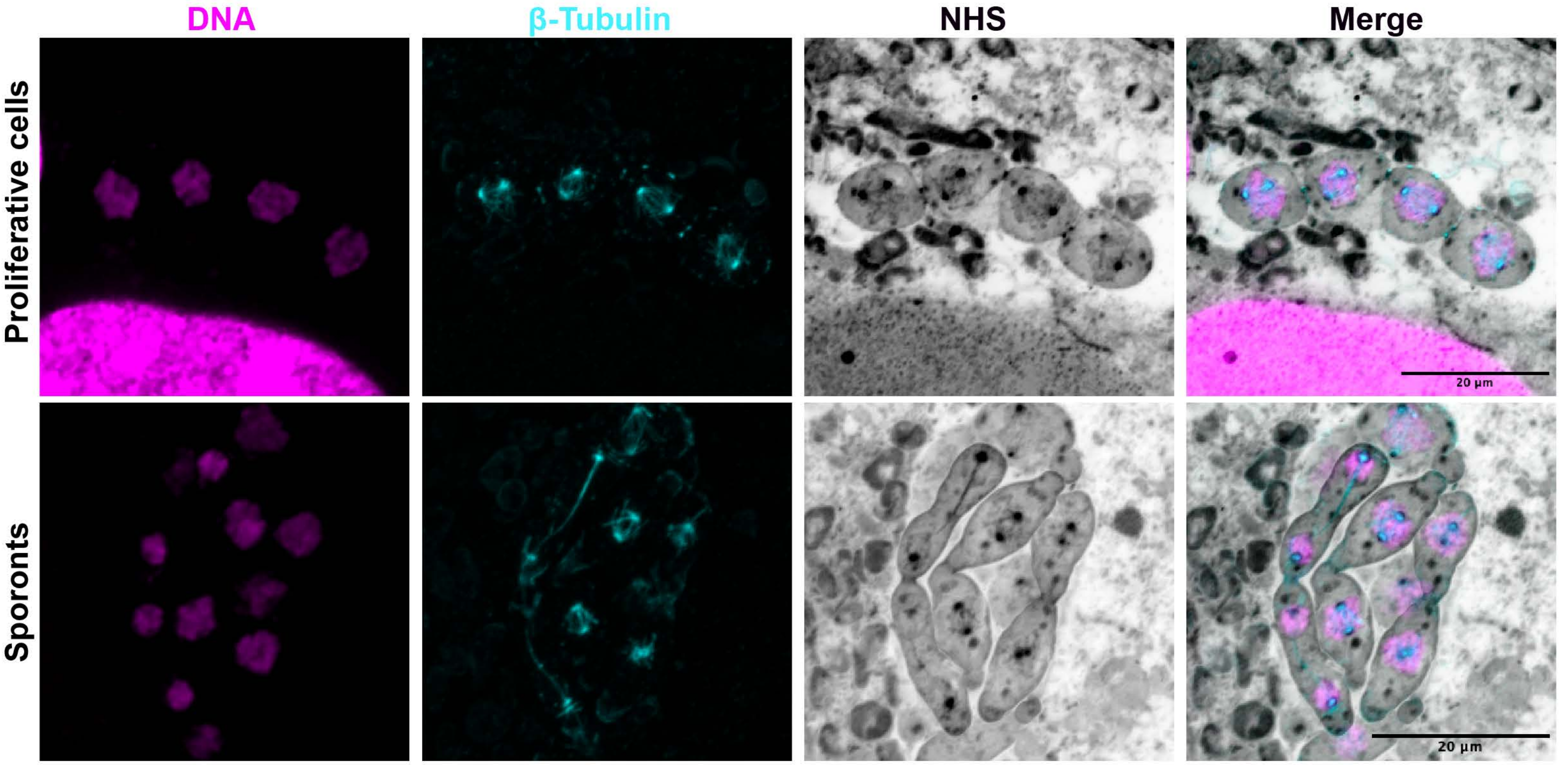


**Figure S4. *E. intestinalis* chains undergo synchronized cell cycle stages.** Representative maximum intensity projection of expanded vero cells infected with *E. intestinalis.* Samples were stained for DNA (DRAQ5, magenta), proteins (NHS-ester, grayscale), and *E. intestinalis* β-Tubulin (custom antibody, cyan). Parasites in the same chains are synchronized at the same cell cycle stages, based on the appearance of β-tubulin. Across three independent experiments, 417 chains were analyzed.

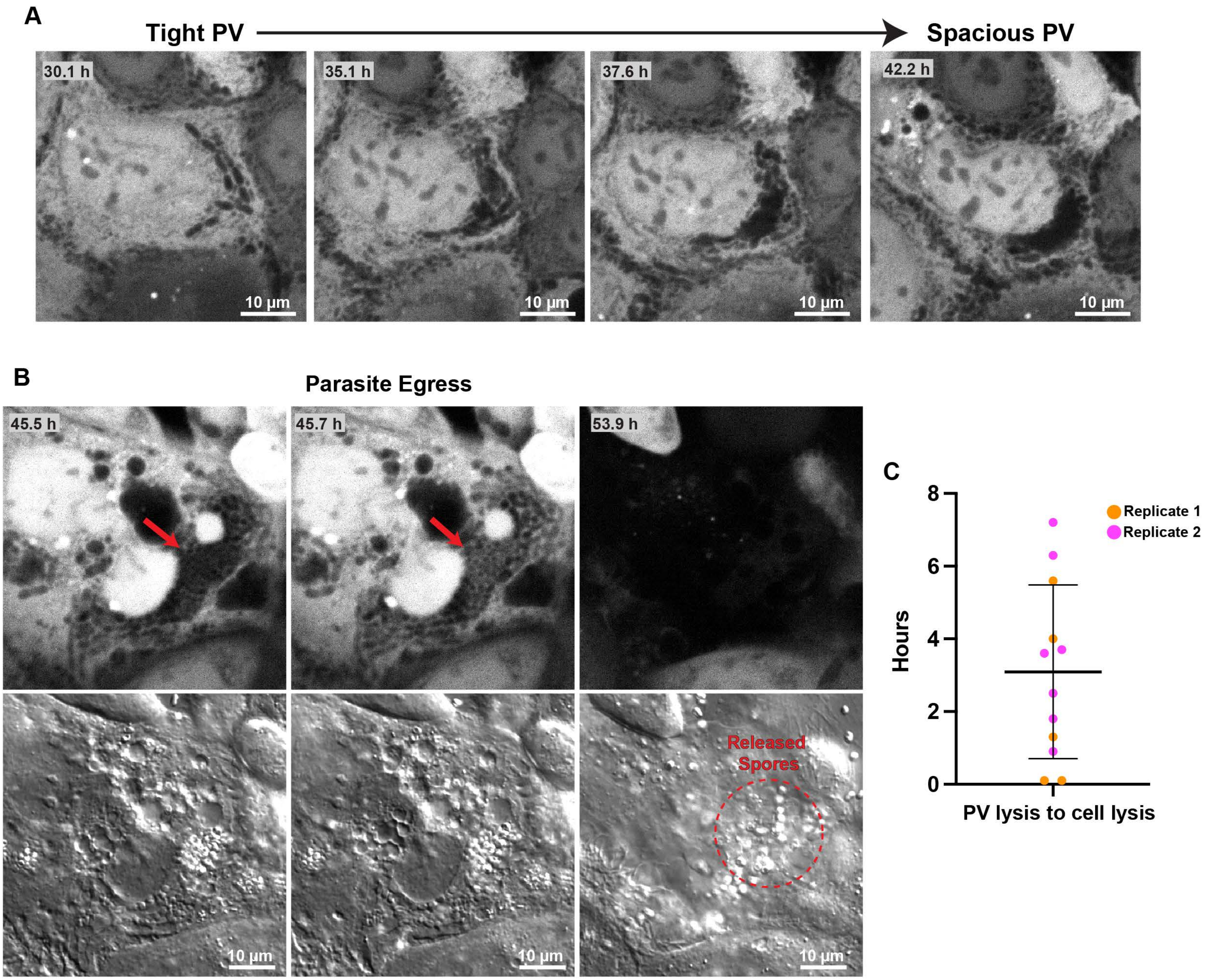


**Figure S5.** ***E. intestinalis*** **egress in Caco-2 cells. A.** Movie frames of *E. intestinalis* infection in Caco-2 cells from 30.1 h h to 42.2 h, showing the transition from a tight PV to a spacious PV using silhouette microscopy. **B.** Silhouette microscopy and corresponding DIC images from movie frames of *E. intestinalis*-infected Caco-2 cells. Timepoints are later in infection, showing PV permeabilization, followed by host cell lysis and parasite egress. All fluorescent images are representative minimum intensity projections. **C.** Quantification of the timing from PV lysis to host cell lysis. Three independent experiments were performed, and the error bar represents SD across the experiments. Replicate 1, n = 5 PVs; Replicate 2, n = 7 PVs; Replicate 3 did not capture any parasite egress events.

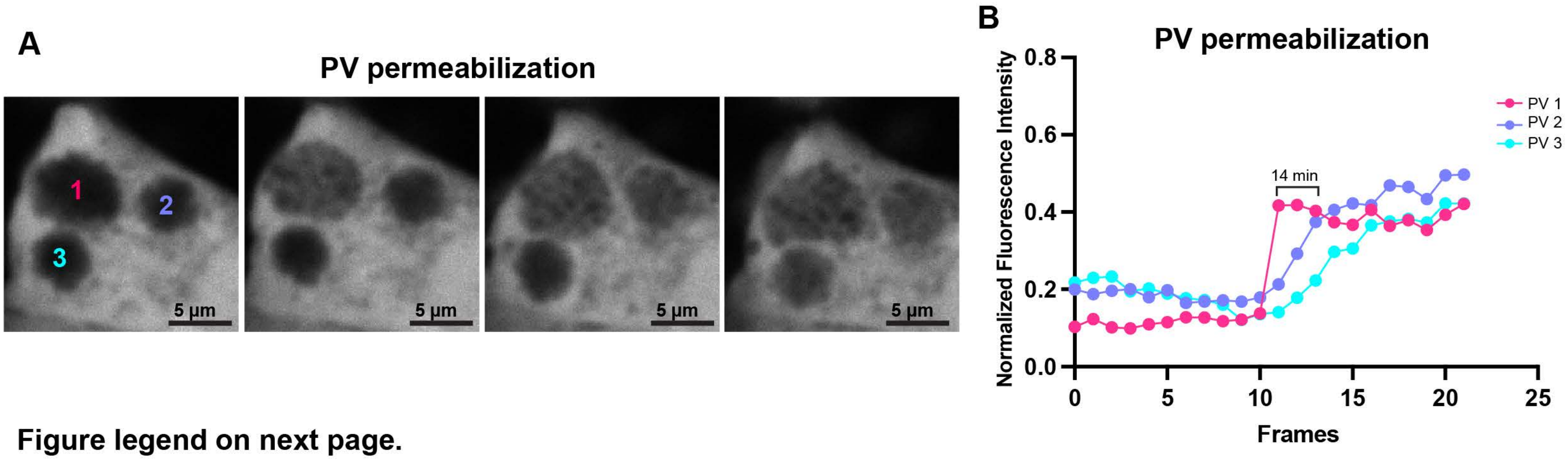


Figure legend on next page.

**C**

**Fusion Events**

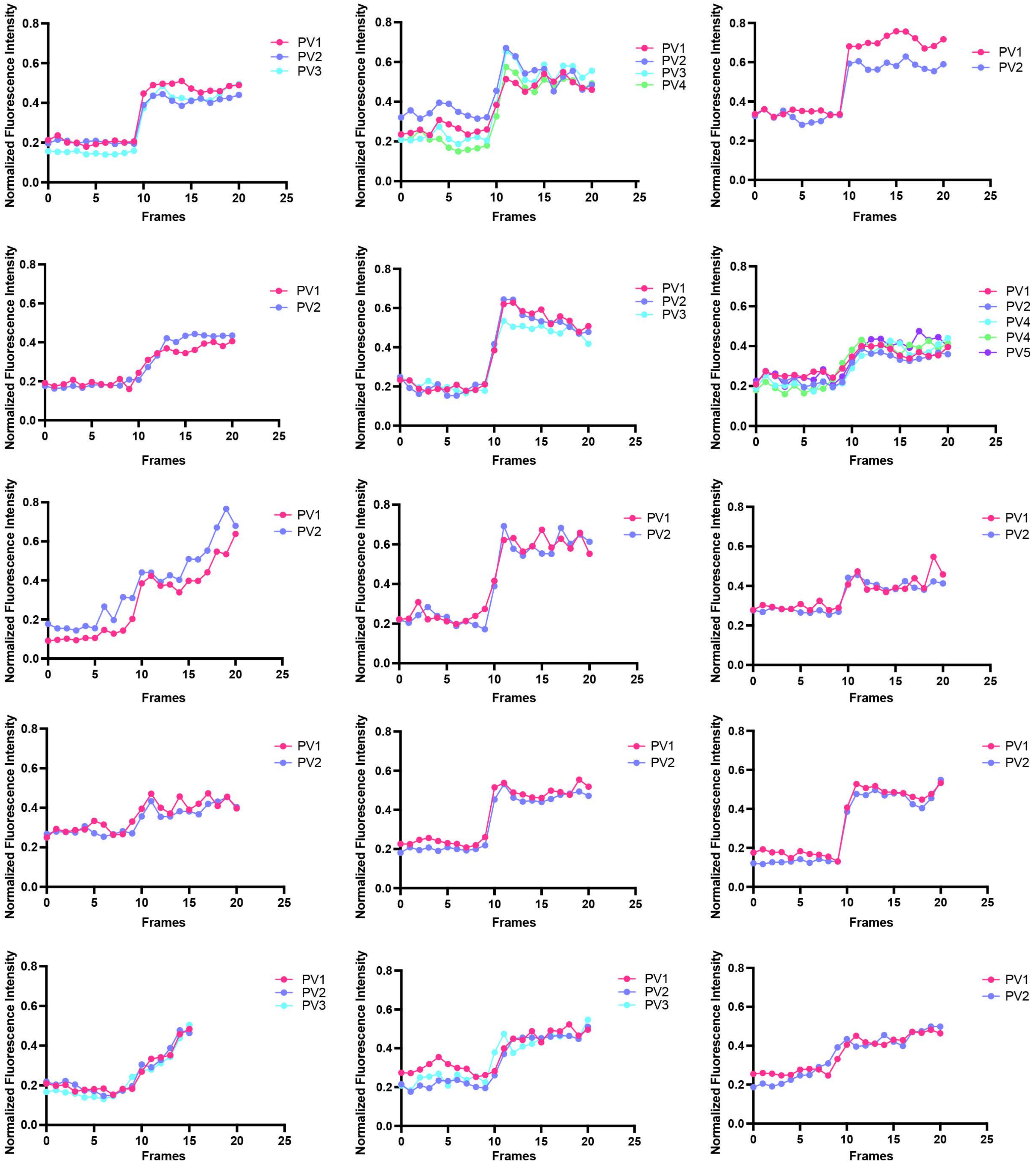


**Figure S6. Fluorescence intensity analysis of PV permeabilization to assess PV fusion. A.** Representative images of PVs that are in close proximity but do not fuse. The PVs clearly rupture at different times, leading to cytoplasmic fluorescence influx at different times. **B.** Quantification of fluorescence intensity of the three PVs in (A). **C.** Fluorescence intensity of 25 examples of fused PVs, measured 10 frames before and after PV permeabilization.

**Dataset S1.** Replication kinetics and PV dynamics for all analyzed live-cell imaging in Vero cells. (Excel spreadsheet)
**Dataset S2.** Replication kinetics and PV dynamics for all analyzed live-cell imaging in Caco-2 cells. (Excel spreadsheet)

### Supplementary Movie Legends

**Movie S1.** Identifying *E. intestinalis* infected Vero cells. Representative silhouette microscopy and DIC movies associated with Figure 1D, played in reverse from a clearly infected later stage back to the original sporoplasm. Parasites are marked by orange outlines.

**Movie S2.** Replication kinetics of *E. intestinalis* in Vero cells. Representative silhouette microscopy and DIC movies associated with Figure 2A. Parasites are marked by orange outlines.

**Movie S3.** Replication kinetics of *E. intestinalis* in Caco-2 cells. Representative silhouette microscopy and DIC movies associated with Figure 2F. Parasites are marked by orange outlines.

**Movie S4.** 3D projection of a sporoblast associated with Figure 5A.

**Movie S5.** *E. intestinalis* infected Vero cells showing the transition from a tight PV to a spacious PV. Representative silhouette microscopy movie associated with Figure 6A. Parasites are marked by orange outlines.

**Movie S6.** *E. intestinalis* infected Caco-2 cells showing the transition from a tight PV to a spacious PV. Representative silhouette microscopy movie associated with Figure S4A. Parasites are marked by orange outlines.

**Movie S7.** *E. intestinalis* infected Vero cells showing PV fusion, followed by permeabilization. Representative silhouette microscopy and DIC movies associated with Figure 6B.

**Movie S8.** *E. intestinalis* infected Vero cells showing PV fission. Representative silhouette microscopy and DIC movies associated with Figure 6C. The PV of interest is marked by an orange outline.

**Movie S9.** *E. intestinalis* exiting Vero cells via host cell lysis. Representative silhouette microscopy and DIC movies associated with Figure 6E. The cell of interest is marked by an orange outline.

**Movie S10.** *E. intestinalis* exiting Caco-2 cells via host cell lysis. Representative silhouette microscopy and DIC movies associated with Figure S4B. The cell of interest is marked by an orange outline.
